# Comparative Genomics Reveals the Ancestral Recombination Landscape of Placental Mammals

**DOI:** 10.64898/2026.04.02.716207

**Authors:** Isabella R. Childers, Nicole M. Foley, Kevin R. Bredemeyer, William J. Murphy

## Abstract

Meiotic recombination is a crucial biological process that ensures proper chromosomal pairing and promotes adaptation. In placental mammals, recombination rates vary widely across species, populations, sexes, individuals, and chromosomes. While the placental X chromosome shows remarkable conservation of both gene order and the recombination landscape across deep evolutionary history, it is unknown whether similar levels of autosomal conservation persist despite extensive chromosomal evolution. Here, we reconstructed an ancestral placental mammal karyotype from chromosome-level assemblies, using slow rates of karyotypic evolution, and inferred an ancestral autosomal recombination map. Analysis of phylogenetic branch lengths and PhyloP-based scores of evolutionary constraint reveals that conserved autosomal regions with low recombination rates have evolved under stronger purifying selection, whereas regions with conserved high recombination rates are less constrained and freer to evolve. Ancestral autosomal regions with low recombination rates were enriched for pathways and GO terms related to cellular function, whereas ancestral regions with high recombination rates were enriched for regulation and some immune-related systems. Tracking the fate of these conserved ancestral recombination hotspots and coldspots across 13 mammal lineages with variable rates of karyotype evolution revealed the retention of autosomal AHRs, but the absence of autosomal ALR conservation. Collectively, our findings reveal variable levels of evolutionary constraint at meiotic recombination in relation to karyotypic evolution, providing new insights into how natural selection influences the evolution of chromosomal organization.

## Introduction

Chromosome evolution in mammals has traversed many paths (Graphodatsky et al. 2011; Dunnum et al. 2001). Extant karyotypes exhibit both diminutive and large diploid chromosome numbers, as shown in the Indian muntjac deer (2n=6,7) and the Bolivian Bamboo Rat (2n=118). Some karyotypes are relatively unchanged from the ancestral state, e.g., sloths, cats, and whales, while others have become highly derived, e.g., mice and shrews. Within clades, karyotypes can be highly conserved (e.g., *Myotis bats,* 2n=44) or highly variable (e.g., *Pipistrellus* bats, 2n=18-52) (Lan et al. 2026). Although the pace of karyotype evolution varies, certain syntenic blocks are conserved across most mammals. For instance, chromosome painting with human-chromosome-specific probes has shown that human chromosomes 3 and 21 are associated (i.e., fused) in the majority of non-primate placental mammal karyotypes across the four major superordinal clades, suggesting that this syntenic association was present in the ancestral placental mammal (Ferguson-Smith and Trifonov 2007). Likewise, the X chromosome of placental mammals has been found to be remarkably conserved in karyotype, sequence, gene content, and gene order (Raudsepp et al. 2004; Delgado et al. 2009; Brashear et al. 2021). More recently, attention has focused on the conservation of the meiotic recombination rate landscape across the mammalian X chromosome, where conserved, collinear boundaries of elevated and reduced recombination rates appear to be present (Li et al. 2019; Foley et al. 2026). The largest recombination deserts tend to preserve the speciation history in clades where hybridization is rampant, which has sparked interest in resolving recombination landscapes to complement phylogenomic analyses (Foley et al. 2026).

Meiotic recombination is an essential process in sexual organisms that enables proper segregation during meiosis and promotes genetic diversity. During prophase I, recombination begins with double-stranded breaks and is resolved as either a “crossover” (i.e., reciprocal exchange between homologous chromosomes) or a “non-crossover” (i.e., resolution of double-stranded breaks through gene conversion; Baudat et al. 2013). Recombination rate is not uniform across the genome; certain loci, denoted “hotspots,” exhibit elevated recombination compared to the rest of the genome (Steinmetz et al. 1982; Chakravarti et al. 1986; Paigen and Petkov 2010). It was later discovered that, in many mammals, a zinc-finger protein, PRDM9, was responsible for positioning recombination hotspots (Baudat et al. 2010; Parvanov et al. 2010; Paigen and Petkov 2018). However, some mammals lack PRDM9 yet still possess recombination hotspots (i.e., canids; Axelsson et al. 2012; Joseph et al. 2024).

While a conserved, undulating landscape of recombination hot and cold spots is present on the X chromosome of many placental mammals (Foley et al. 2026), recombination rates on autosomes are expected to vary across species, populations, sexes, individuals, and along chromosomes (Peñalba and Wolf 2020). Despite this variability, some common trends have been observed. Longer chromosomes tend to have higher recombination rates toward the ends and lower rates near the middle, while smaller chromosomes usually exhibit higher recombination rates across most of their length (Haenel et al. 2018). However, there is limited prior literature on the evolutionary conservation of autosomal recombination rates in diverse clades. A few studies have shown that immunity genes are common in recombination hotspots within specific lineages, an adaptive strategy that creates allelic diversity through admixture (Vandiedonck and Knight 2009; Foley et al. 2024). Finding these conserved regions at the extremes of recombinational variation may offer a unique way to measure the functional coherence and constraints on gene clustering and long-range synteny conservation.

Here, we extended our previous observations of the conserved X-chromosome landscape by comparing the autosomal recombination landscape across divergent placental mammals to identify additional genomic regions with conserved recombinational properties. To address this question, we used a novel approach to reconstruct the ancestral karyotype of placental mammals and infer the ancestral recombination landscape, selecting mammals previously identified as having slowly evolving karyotypes. We predicted that this approach would reduce phylogenetic noise in our ancestral reconstruction caused by excessive lineage-specific rearrangements. We generated new long-read-based genome assemblies for the aardvark and Hoffmann’s two-toed sloth and used the machine-learning algorithm ReLERNN (Adrion et al. 2020) to estimate recombination maps for both from population genomic data. From this ancestral reconstruction, we traced the stability and functional properties of the recombination landscape in descendant lineages that have evolved with variable recombination rates. We used these results to infer the functional and selective constraints on synteny evolution relative to recombination over the past 100 million years of placental mammal evolution.

## Results

### Chromosomal-level assemblies of slowly evolving karyotypes in Afrotheria and Xenarthra

To facilitate our reconstruction of the ancestral placental mammal genome, we sequenced chromosome-level genome assemblies of the aardvark and the sloth, representatives from Afrotheria and Xenarthra, which exhibit slow rates of karyotypic evolution (Svartman et al. 2006; Yang et al. 2003) (**Table 1**). We generated PacBio CLR and Illumina reads for a male individual from each species. PacBio reads were sequenced to 59x and 68x coverage, and Illumina reads were sequenced to 132x and 82x coverage for the aardvark and sloth, respectively. Consistent with previously published chromosome painting data, our primary genome assemblies for the sloth and aardvark contained 25 and 10 chromosomes, respectively. However, we were unable to assemble a Y chromosome for either individual, as the Hi-C data used for scaffolding were derived from unrelated female specimens. The final primary assembly length is 4.2 Gb for the aardvark (N50 = 386 Mb) and 3.1 Gb for the sloth (N50 = 153 Mb). The accuracy of both genomes was high, with QV values of 31.9 for the aardvark genome assembly and 37 for the sloth genome assembly. Compleasm, which reports the percentage of single-copy genes in a genome assembly, further indicated that our assemblies were of high quality, with 91.3% of genes in the aardvark and 93.7% in the sloth.

**Table 1.**
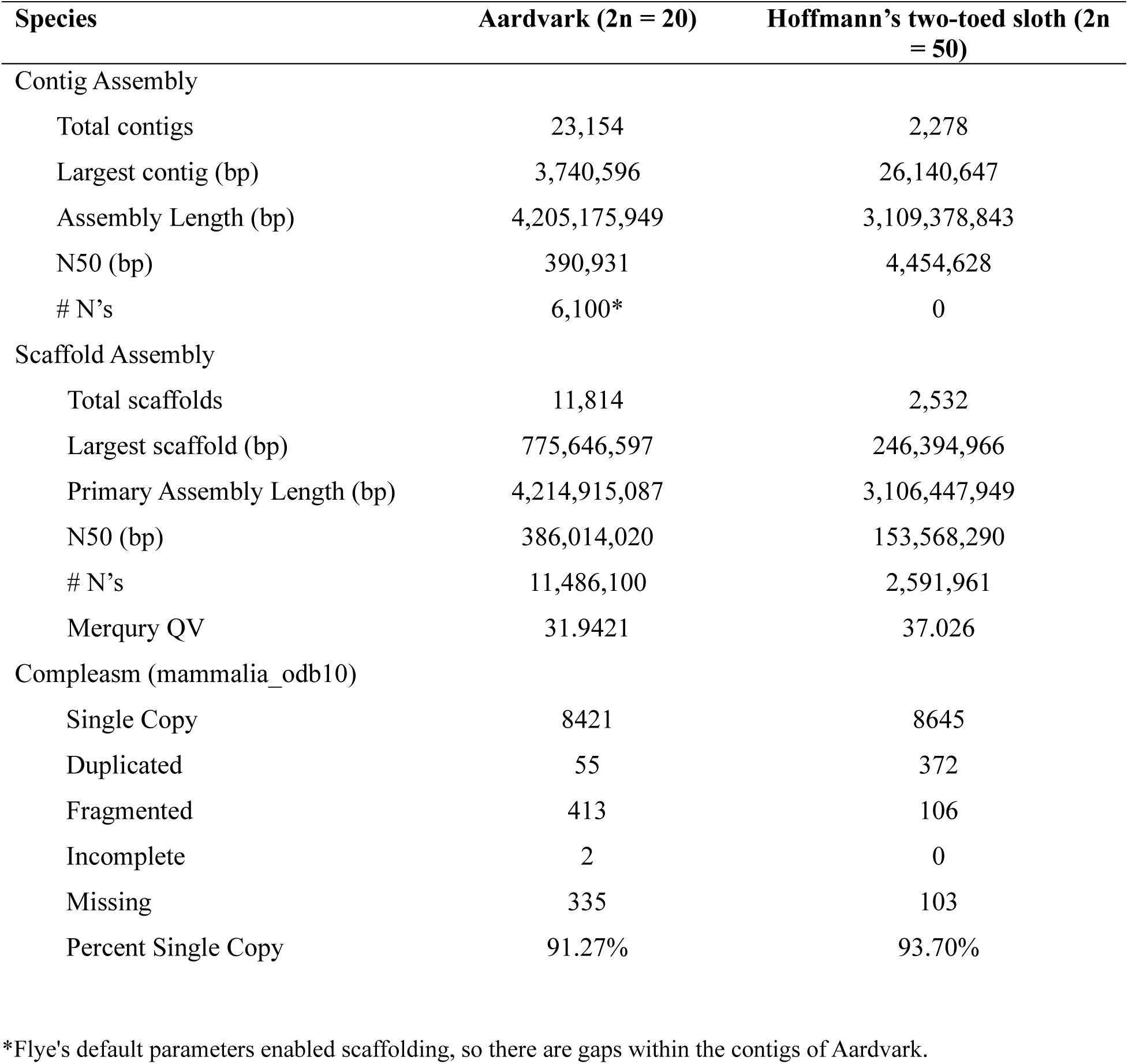
Aardvark and Hoffmann’s two-toed sloth genome assembly statistics.

| **Species** | **Aardvark (2n = 20)** | **Hoffmann's two-toed sloth (2n = 50)** |
| --- | --- | --- |
| **Contig Assembly** |  |  |
| Total contigs | 23,154 | 2,278 |
| Largest contig (bp) | 3,740,596 | 26,140,647 |
| Assembly Length (bp) | 4,205,175,949 | 3,109,378,843 |
| N50 (bp) | 390,931 | 4,454,628 |
| # N's | 6,100* | 0 |
| **Scaffold Assembly** |  |  |
| Total scaffolds | 11,814 | 2,532 |
| Largest scaffold (bp) | 775,646,597 | 246,394,966 |
| Primary Assembly Length (bp) | 4,214,915,087 | 3,106,447,949 |
| N50 (bp) | 386,014,020 | 153,568,290 |
| # N's | 11,486,100 | 2,591,961 |
| Mercury QV | 31.9421 | 37.026 |
| **Compleasm (mammalia_odb10)** |  |  |
| Single Copy | 8421 | 8645 |
| Duplicated | 55 | 372 |
| Fragmented | 413 | 106 |
| Incomplete | 2 | 0 |
| Missing | 335 | 103 |
| Percent Single Copy | 91.27% | 93.70% |

\*Flye's default parameters enabled scaffolding, so there are gaps within the contigs of Aardvark.

### Reconstructing the Recombination Landscape in the Ancestral Placental Mammal Genome

The ancestral placental mammal genome was reconstructed using chromosome-level assemblies from extant mammals previously identified as having slowly evolving karyotypes. We sampled placental mammals that included at least one representative from each of the four main superordinal groups (Murphy et al. 2021): Xenarthra (Hoffmann’s two-toed sloth), Afrotheria (aardvark), Laurasiatheria (the blue whale and domestic cat), and Euarchontoglires (human). By comparing protein-coding genes and their alignments, we observed a high level of synteny and conservation of gene order among these five mammals, with few rearrangements (**Fig. 1A**). This result was consistent with previous Zoo-FISH studies indicating that these taxa are very close to the ancestral placental mammal karyotype (Svartman et al. 2012). We identified many common human chromosome syntenic associations (HCSA), including 3/21, 4/8p, 14/15, 10/22/12p/22 (**Fig. 1B**), 7b/16p, and 12q/22 (**Table S1**). Human chromosomes 5, 9, 11, 13, 14, 17, 18, 20, 21, and X appeared on one chromosome block in each mammal, while chromosomes 7, 12, 16, and 19 were found on two blocks (**Table S1**). The remaining human chromosomes—1, 2, 3, 4, 6, 8, 10, 15, and 22—were present on one or a few chromosome blocks (**Table S1**).

**Figure 1.**
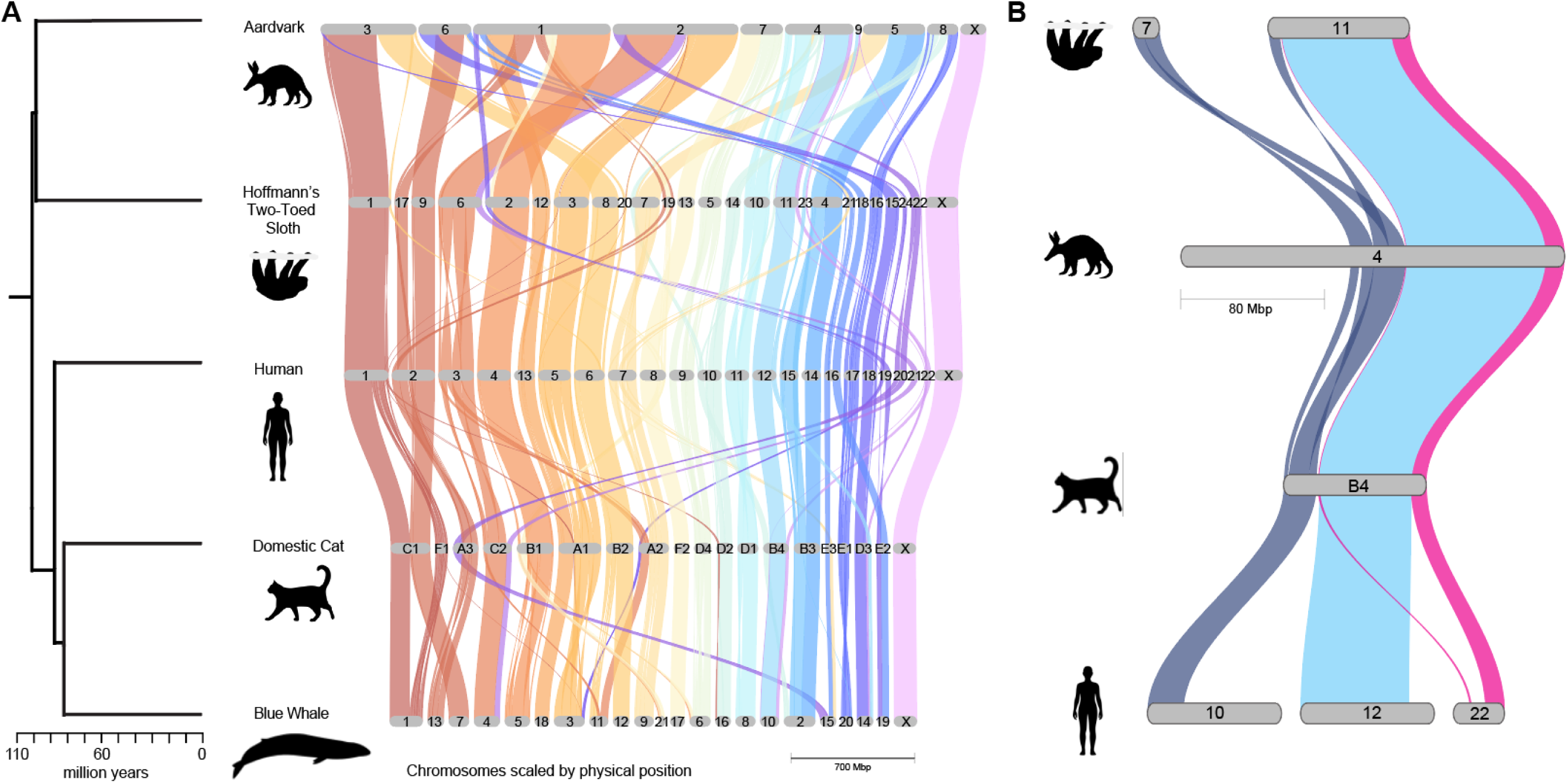
Extensive conservation of collinearity across five ordinal lineages of placental mammals identifies large syntenic blocks likely present in the ancestor. The gene synteny plot was generated by GENESPACE. Solid bands indicate regions of gene collinearity shared among the five species, and twisted bands represent inversions or rearrangements. Colored bands connecting chromosomes indicate conserved synteny relative to the human. A) Collinearity is shown between the five mammals with slowly evolving karyotypes. The left phylogeny shows the timetree of the five placental mammals, derived from Murphy et al. (2021). B) Example of chromosome association of human chromosome 10+22+12+22 in domestic cat, aardvark, and sloth. Credit: Silhouettes were reproduced from PhyloPic (https://www.phylopic.org/) under a CC0 1.0 Universal Public Domain license unless otherwise stated. sloth, created by Pearson Scott Foresman; human, created by Cagri Cevrim; cat, created by Skye McDavid, and whale, created by Chris Huh, under an Attribution-Share Alike 3.0 Unported license https://creativecommons.org/licenses/by-sa/3.0/. Alt text: Graphical representation of collinearity across divergent mammals. The left figure shows synteny among the five species used in the ancestor reconstruction with their phylogenetic tree. The right figure shows a similar collinearity plot, but highlighting an example of a chromosome association in three mammals (i.e., human chromosomes 10, 22, 12, and 22).

The DESCHRAMBLER algorithm (Kim et al. 2017) reconstructed an ancestral karyotype containing 28 ancestral predicted chromosome fragments (APCFs) (**Table S1**). In our reconstruction, five ancestral chromosomes partially agree with Svartman (2012) (**Table S1**). Human chromosome 17 appears in two APCFs, although in Svartman (2012)’s Zoo-FISH-based reconstruction, it is found on a single ancestral chromosome. Similarly, human chromosome 10q appears in two separate APCFs, whereas Svartman (2012) places it on a single ancestral chromosome. Human chromosomes 10, 12, and 22 are located on one ancestral chromosome in Svartman (2012), whereas DESCHRAMBLER separates chromosomes 12 and 22 from chromosome 10 (**Table S1**). Additionally, Svartman (2012) reports that smaller fragments from chromosomes 12 and 22 are on a second single ancestral chromosome, yet DESCHRAMBLER separates chromosome 12 from 22 (**Table S1**). Finally, human chromosome 15 is found on a different APCF from the APCF associated with 14/15 (**Table S1**); however, in Svartman (2012), it is in a single chromosome. We also varied the block size, outgroup, and reference choice and showed that DESCHRAMBLER reconstructions were largely robust to changes in key parameters (**Tables S1 and S6; Supplementary Methods**).

Based on shared syntenic regions among the four non-human mammals, we grouped APCFs into a single ancestral chromosome, yielding a predicted ancestral chromosome number of 23 that is largely consistent with the Zoo-FISH-based ancestors (**Fig. 2**). Our results clarify several previously controversial patterns. First, our results provide support for the presence of the human 10+12+22 association in the ancestral placental mammal karyotype. Although it is sporadically present across the phylogeny, it does not appear to have independently evolved in the carnivores (cat), sloth, and aardvark as evidenced by small homologous adjacent regions of 10+22+12+22 shared in each of these three species that were previously undetectable by Zoo-FISH (**Fig. 1B**). Second, our whole genome alignments that include a new aardvark and two-toed sloth genome assemblies confirm that human chromosome 1 is highly collinear across its length in Afrotheria (aardvark), Xenarthra (sloth), and Laurasiatheria (cat and blue whale) (**Fig. 1**), and likely represents a retention of the ancestral conformation (Murphy et al. 2003).

**Figure 2.**
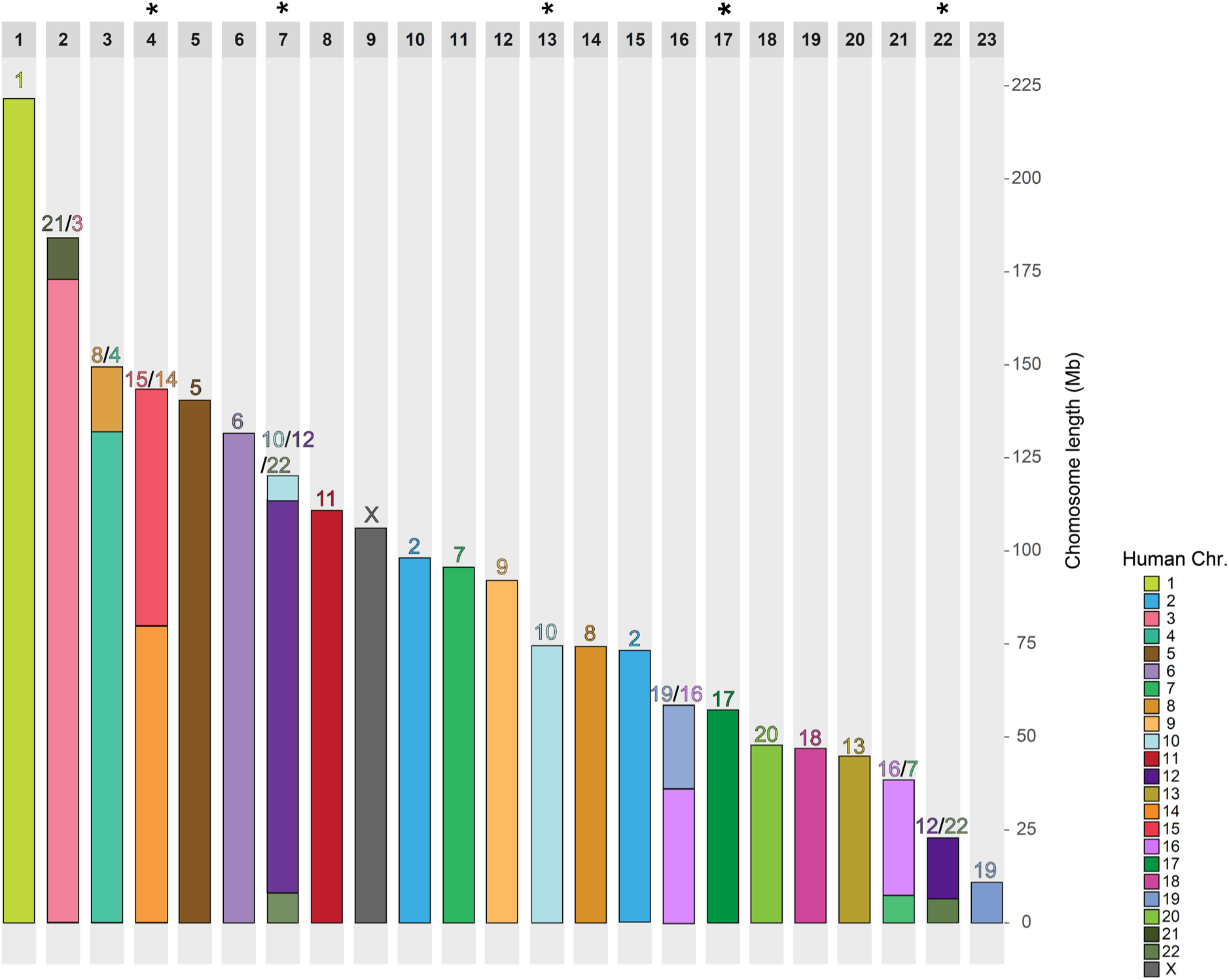
Chromosomal ideogram of the reconstructed ancestral placental mammal karyotype with human synteny. Each column represents one ancestral chromosome with the length in Mb (x-axis). Colored blocks indicate collinearity between the human and ancestral chromosomes. Asterisks mark where two chromosome fragments from the DESCHRAMBLER reconstruction were fused to match the predicted ancestral chromosomes hypothesized based on chromosome painting (Svartman 2012). Alt text: Graphical representation of the reconstructed ancestral placental mammal karyotype. Each ancestral chromosome is shown vertically with numbers above and colors representing the human chromosome association. Five asterisks are above ancestral chromosomes 4, 7, 13, 17, and 22.

To infer the ancestral recombination landscape, we used either published data (human and cat; Halldorsson et al., 2019; Foley et al., 2026) or newly constructed recombination maps (aardvark and sloth) generated by applying the machine-learning algorithm ReLERNN (Adrion et al., 2020) to population-genomic data. The resulting recombination landscapes for aardvark and two-toed sloth exhibit features predicted by their chromosome lengths and sizes. In long chromosomes, higher recombination rates tend to occur at the ends, while lower rates are seen toward the middle (e.g., aardvark chromosomes 1 and 2 in **Fig. S3**). Short chromosomes display elevated recombination rates throughout their length (e.g., aardvark chromosome 9 and sloth chromosome 19 in **Figs. S3 and S4**).

The recombination map, representative of each species, was divided into four recombination rate categories (quartiles) (**Fig. S1-S4**). We identified collinear genomic regions that have consistently remained in high or low recombining regions across all species throughout mammalian evolution (which we infer as our “ancestral recombination landscape”) (**Fig. S5-S8**). In humans, ancestral cold spots were identified in 166 loci across 22 of 23 chromosomes, while hot spots were detected in 177 loci across all 23 chromosomes (**Fig. S5**). In the domestic cat, ancestral cold spots were identified at 163 loci and ancestral hot spots at 176 loci across all 19 chromosomes (**Fig. S6**). In the aardvark, ancestral cold spots were identified in 170 loci and hot spots in 179 loci across all 10 chromosomes (**Fig. S7**). In Hoffmann’s two-toed sloth, ancestral cold spots were detected in 160 loci in 22 of the 24 chromosomes, while hot spots were identified in 179 in 23 of the 24 chromosomes (**Fig. S8**).

Ancestrally low (ALR) and ancestrally high recombining regions (AHR) were identified on syntenic regions of the reconstructed ancestral placental mammal genome (**Fig. 3**). Cold spots in the ancestor were found in 196 loci across 22 of 23 chromosomes, while hot spots were identified in 205 loci across all 23 chromosomes (**Fig. 3**). As expected, most chromosomes in the reconstructed placental mammal ancestor show higher recombination rates near their ends and lower rates toward the middle (e.g., chromosomes 1, 14, and 20 in **Fig. 3**), suggesting these patterns have been conserved for over 100 million years of independent evolution. Smaller human chromosomes, such as 21 and 22, which are situated at the ends of ancestral chromosomes, exhibit exclusively high recombination rates (see ancestral chromosomes 2, 7, and 22 in **Fig. 3**). However, most of the shorter ancestral chromosomes did not possess exclusively elevated recombination rates, except for ancestral chromosome 23 (**Fig. 3**).

**Figure 3.**
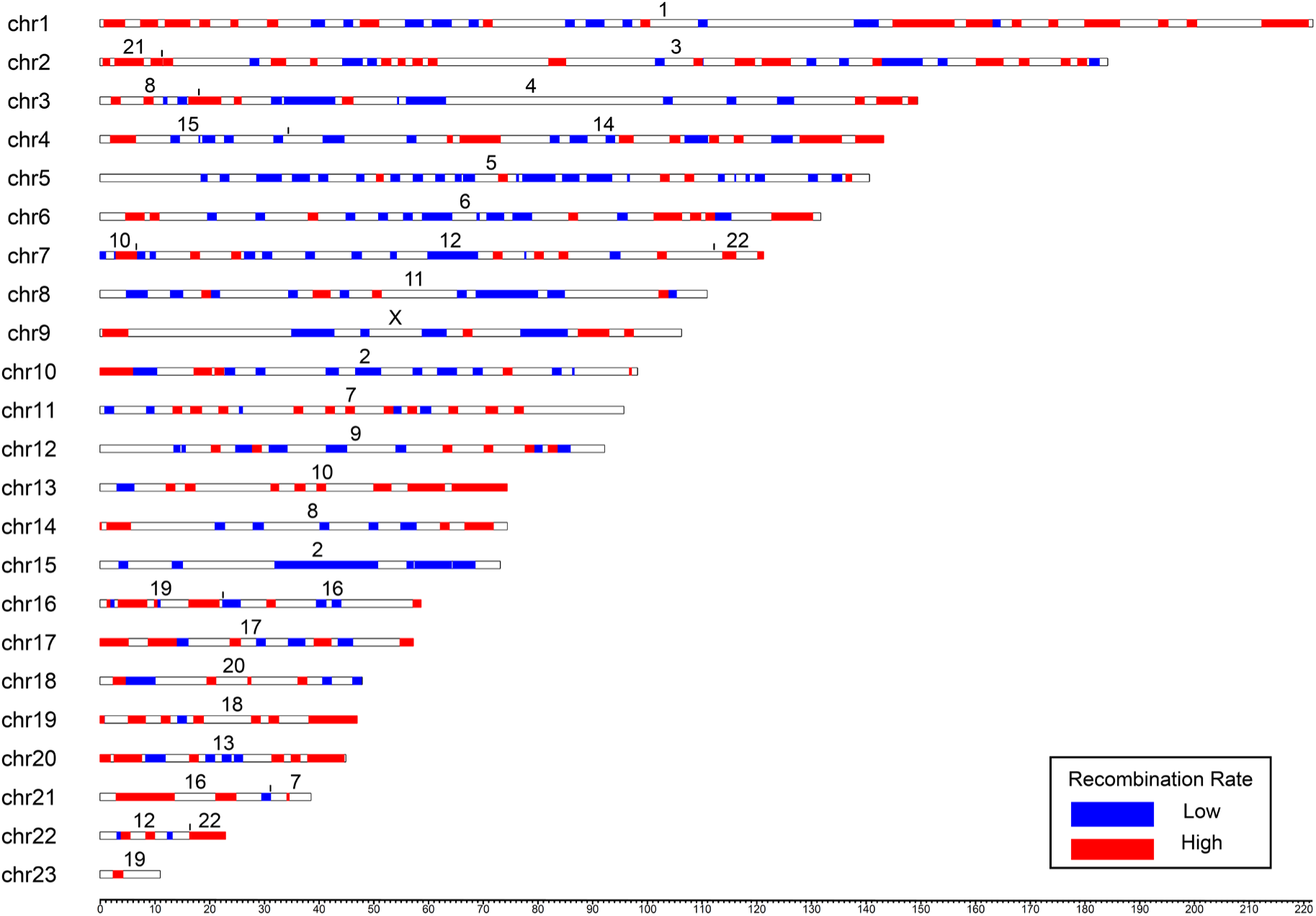
Ideogram of the reconstructed ancestral placental mammal genome depicting syntenic regions where descendant lineages have retained similar low and high meiotic recombination rates across three or more of the species (human, cat, sloth, and aardvark), each representing one of the four placental mammal superorders. We hypothesize these represent the ancestral recombination states in the placental mammal ancestor. Colors along the chromosomes indicate conserved syntenic regions possessing conserved low (blue) and high (red) recombination rates. The numbers above each chromosome indicate synteny relative to a human chromosome. The x-axis depicts the length of the chromosomes in base pairs. Alt text: Graphical representation of the ancestral low and high recombining regions highlighted on an ideogram of the ancestral placental mammal karyotype that was reconstructed in this study.

### Phylogenomic properties and Evolutionary Constraint of ALR and AHR genes in karyotypically diverse mammals

Recombination interacts with natural selection to influence the retention of genomic ancestry along chromosomes (Schumer et al. 2018). When genomic ancestries are inferred using phylogenomic approaches, the distribution of local locus trees that support the species history (versus local gene trees consistent with ILS or introgression) can be reliably predicted from recombination rates (Burbrink et al. 2025). A recent comparative genomic study identified a large, conserved X-chromosome recombination desert shared across placental mammals, enriched for speciation history and resistant to gene flow (Foley et al. 2026). This finding suggests that this genomic region may serve as a robust phylogenomic marker for clades whose relationships are likely obscured by introgression. Therefore, we searched our inferred ancestral autosomal landscapes for large regions with conserved high or low recombination that could also be more useful for phylogenomic studies across mammals by characterizing them with respect to measures of evolutionary constraint based on PhyloP scores from the Zoonomia alignment (Christmas et al. 2023). When we compared the topologies, PhyloP scores, and branch lengths of clades inferred from ALR and AHR regions of the Zoonomia 240 species genome alignment (Zoonomia Consortium 2020; Foley et al. 2023), relationships consistent with the species tree were found in the ALR tree for species known to hybridize, whereas relationships reflective of past introgression were recovered from the AHR tree (**Fig. 4a, S9, S10**) (see Foley et al. 2023). Node heights and branch lengths from the ALR tree inferred from ALR regions were also shorter than those inferred from AHR regions (**Fig. 4b** and **4c**). PhyloP scores were more constrained in ALR regions relative to AHR regions (**Fig. 4d**).

**Figure 4.**
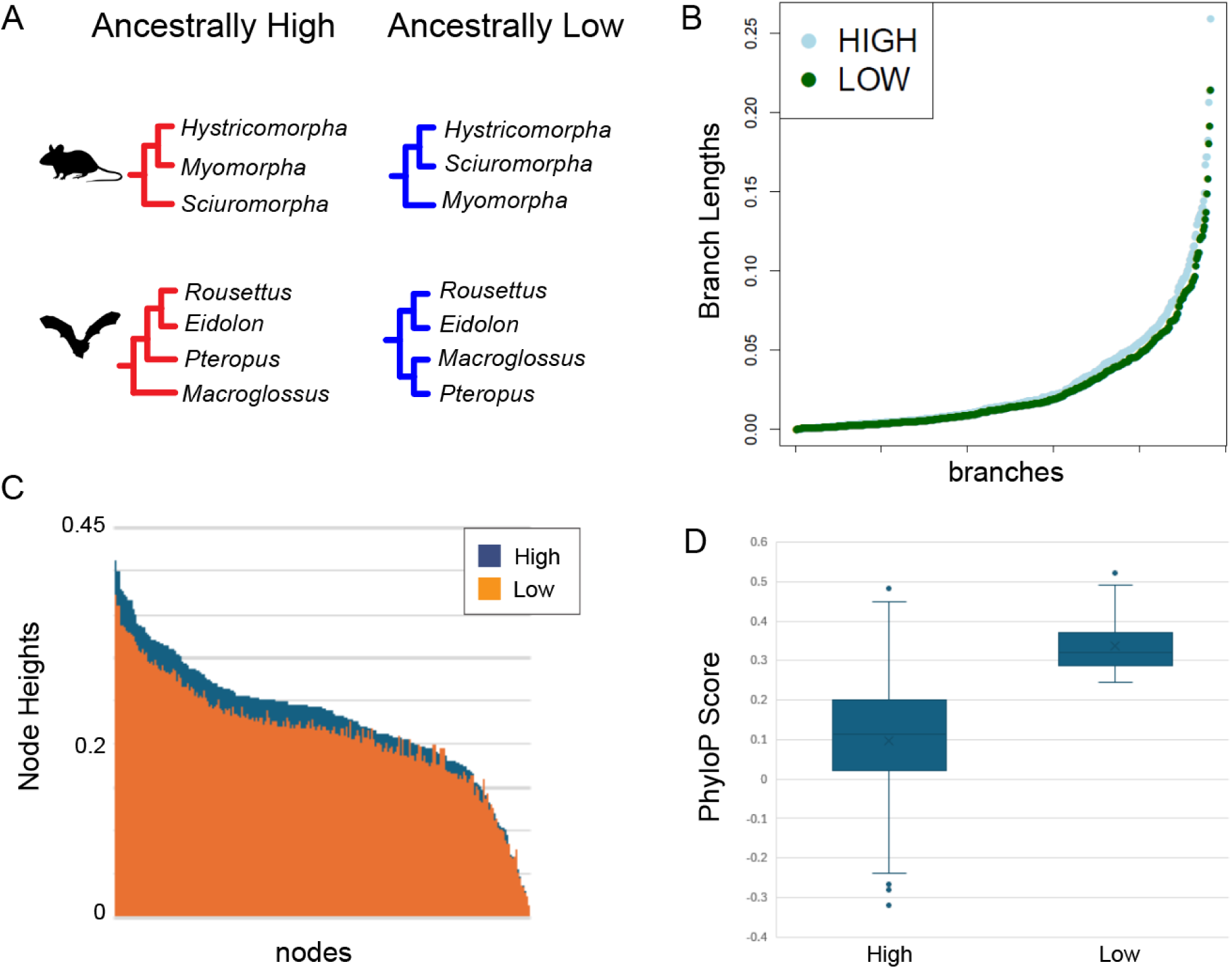
Phylogenomics of ancestrally conserved high- and low-recombining regions. Differences in A) topology, B) branch lengths, and C) node heights between trees derived from ancestrally conserved high and low recombining regions. D) Comparison of PhyloP per-base measures of selective constraint in conserved high and low recombining regions. Credit: Silhouettes were reproduced from PhyloPic (https://www.phylopic.org/) under a CC0 1.0 Universal Public Domain license unless otherwise stated; bat, created by Margot Michaud; mouse, created by Anthony Caravaggi. Alt text: Four figures that depict the phylogenomic analysis of this study. The top-left figure, or figure A, depicts two phylogenetic trees of two different clades sampled from ancestrally high or low regions. The top-right figure, or figure B, depicts the branch lengths between the high- and low-recombining regions, with the high shown as having longer branch lengths. The bottom-left figure, or figure C, shows the node heights between the low and high recombining regions, with the high regions having higher node heights. The bottom-right figure, or figure D, shows the PhyloP scores between high- and low-recombining regions, with low having higher scores and a narrower range than high.

### Functional genetic properties of ALR and AHR genes

Given that PhyloP scores were more conserved in ALR regions, we sought to quantify the relative distribution of genes across key biological pathways. Ideally, this approach would identify genes we hypothesize represent “core conserved” components of a pathway (ALR) and those we hypothesize are freer to vary (AHR). To do this, we first queried the human- and cat-referenced autosomal gene sets from the ancestor-inferred ALR (2,530 genes) and AHR (3,182 genes) regions (**Table S3**) against the databases within WebGestalt (Elizarraras et al. 2024) and STRING (Szklarczyk et al. 2023). Autosomal ALR regions are enriched for a variety of critical cellular processes, including DNA recombination, metabolic pathways, and keratinization (**Fig. 3A**; **Table S4**). Autosomal AHR regions were more enriched for signaling pathways, development, and the regulation of cellular processes (**Fig. 3A**; **Table S4**).

To further investigate the retention of these regions in other genomes, we compared the number of genes in the ALR and AHR human- and cat-reference gene sets that remain in low-and high-recombining regions across representative species from mammal clades characterized by more variable rates of karyotype evolution. These mammals include aardvark (2n = 20), human (2n = 46), Hoffmann’s two-toed sloth (2n = 50), domestic cat (2n = 38), domestic dog (2n = 78), Asian elephant (2n = 56), Nine-banded armadillo (2n = 64), cow (2n = 60), pig (2n = 38), *Myotis* bat (2n = 44), mouse (2n = 40), blue whale (2n = 44), and white rhino (2n = 82) (Foley et al. 2026). We hypothesized that natural selection would constrain certain genes/genomic regions to reside within regions of reduced or elevated recombination (O’Brien et al. 1999). Species with higher diploid numbers generally have more rearranged (derived) karyotypes than those with lower numbers (O’Brien et al. 1999). Chromosome rearrangements should change the relative positions of genes within a chromosome as well as change the overall length in higher diploid numbers (**Fig. S9**). These should disrupt the ancestral recombination rates in those lineages.

We then narrowed our autosomal gene sets to 1,138 in low-recombining regions and 1,565 in high-recombining regions, reflecting genes annotated across all 13 mammals. A hallmark pathways analysis (Liberzon et al., 2015) of these genes revealed that AHR regions are skewed towards pathways involved in UV response to DNA damage, inflammatory response, and estrogen response (**Fig. 5B**). ALR regions are skewed towards more conserved pathways, including DNA repair, unfolded protein response, and TGF beta signaling. Each mammal retained at least 45% of genes in the autosomal ALR and AHR regions (**Fig. 6A**). There were 0 genes from the ALR gene set that remained in the lowest recombination quartile across all 13 mammals (**Table S5**). On the other hand, 17 genes from the AHR gene set remained high across all 13 mammals (**Table S5**). Those genes include *CDC42EP4*, *COG1*, *CPSF4L*, *CPXM2*, *EFR3A*, *FAM104A*, *HTR6*, *NBL1*, *OTUD3*, *PAX7*, *RNF186*, *SDK2*, *SSTR2*, *TMCO4*, *UBR4*, *VTI1A*, and *ZFAT*.

**Figure 5.**
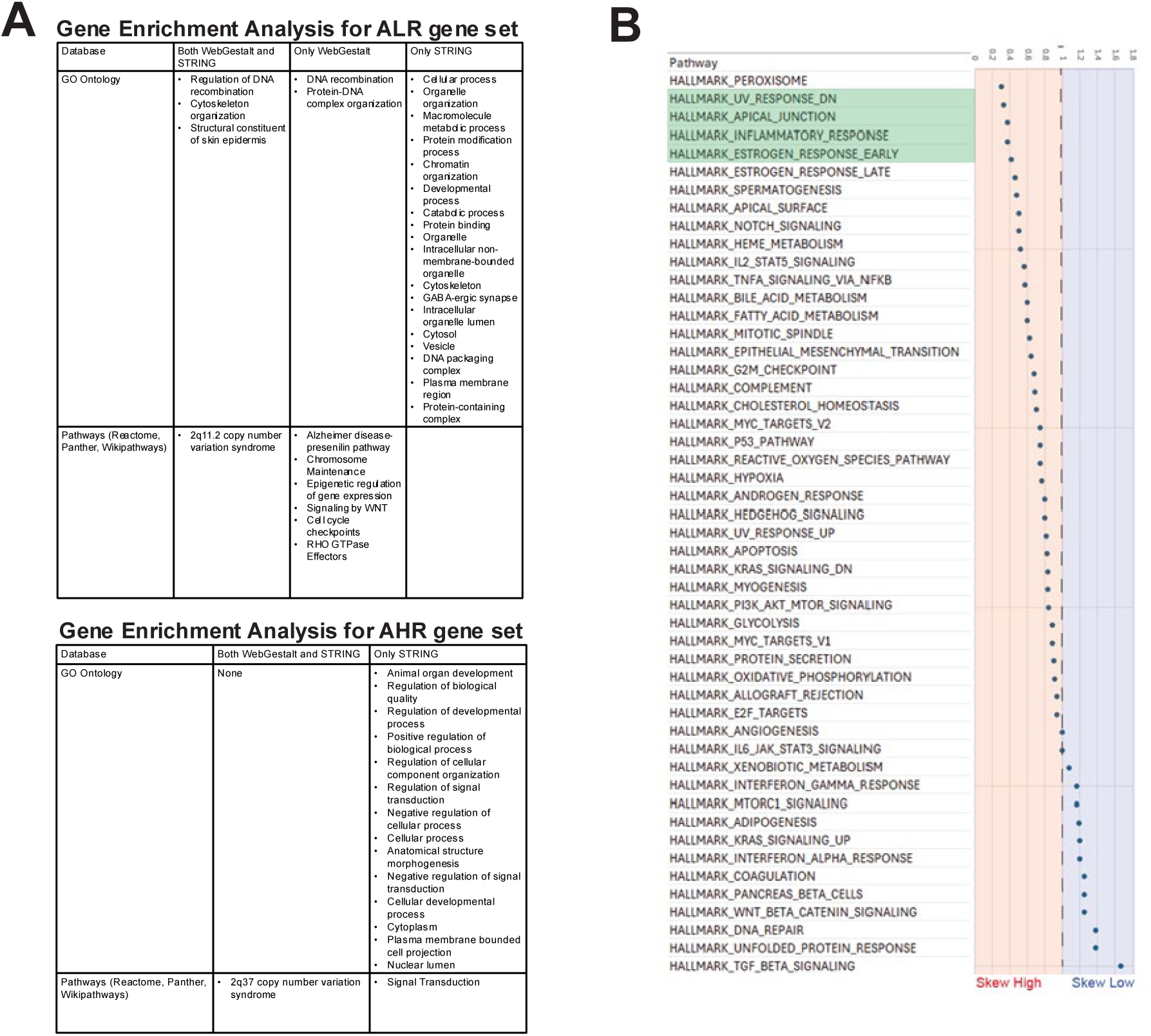
Functional gene enrichment analysis of ALR and AHR. A) Gene enrichment analysis for ALR and AHR autosomal human- and cat-referenced gene sets. GO Ontology and pathway databases are highlighted with terms from both WebGestalt and STRING, from WebGestalt only, or from STRING only. To avoid redundancy, these tables show terms after applying weighted set cover for WebGestalt and merging rows with similarity >= 0.5 in STRING. B) Distribution of genes found in the 50 Hallmark Pathways with respect to ancestrally conserved regions of recombination. A ratio of 1 indicates that genes associated with a given pathway are equally distributed between high- and low-recombination regions. The majority of pathways have more genes in high-recombining regions than in low-recombining regions. Pathways highlighted in green show a significant skew toward high-recombining regions after correcting for the total number of genes in the pathway. Alt text: The left figure shows a table highlighting the gene enrichment analysis for ALR and AHR gene sets from the GO ontology and pathway databases. The right figures show the distribution of genes across the 50 hallmark pathways, skewed towards high- or low-recombining regions.

**Figure 6.**
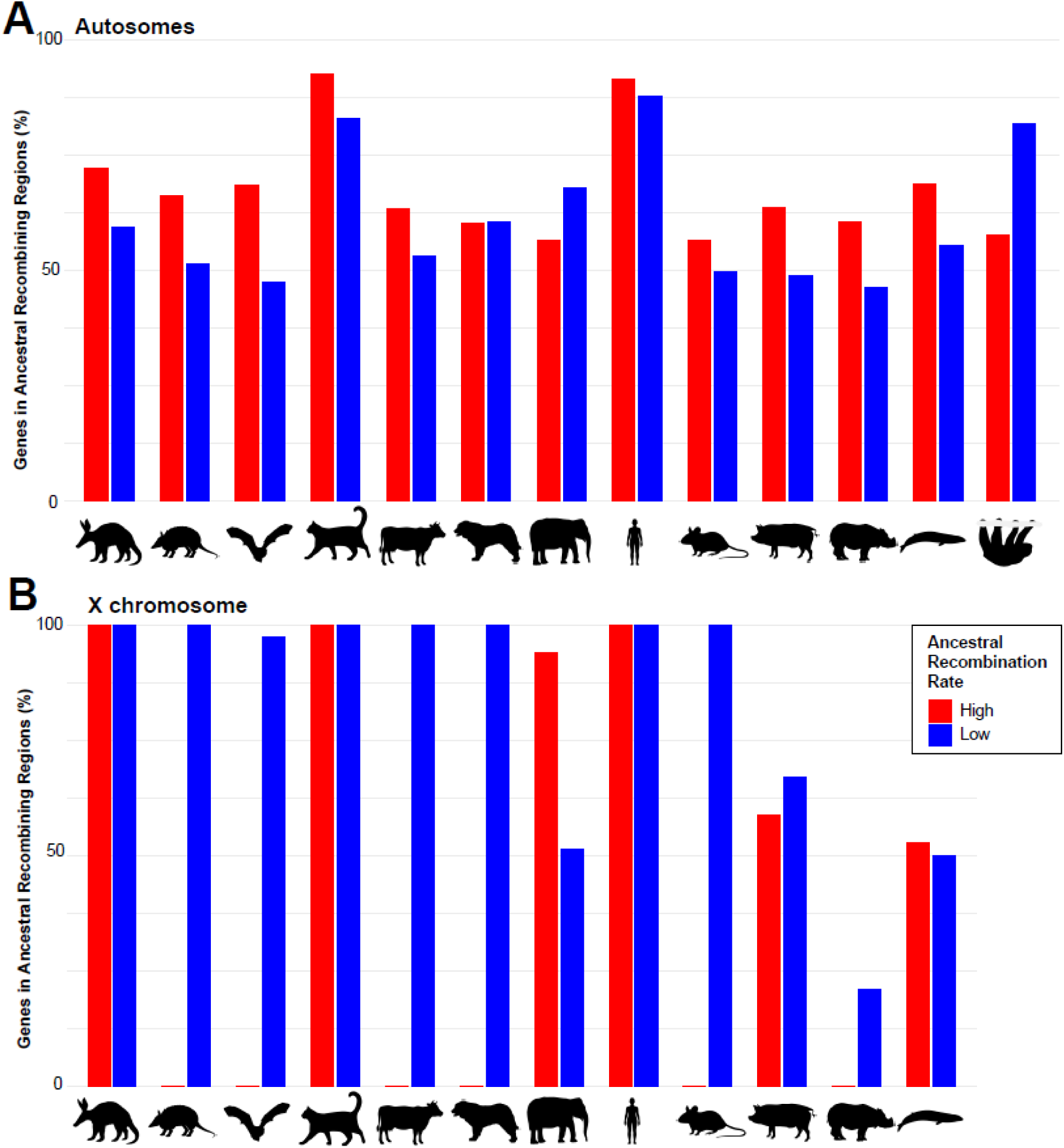
Percentage of genes in ALR and AHR regions that have remained in low and high-recombining regions in mammals with variable karyotype evolution. A) Bar graph depicts the percentage (y-axis) of the autosomal ALR (labeled L and blue) and AHR (labeled H and red) gene set in each mammal’s low and high recombining regions (x-axis). B) Bar graphs depict the percentage (y-axis) of the X chromosomal ALR (labeled L and blue) and AHR (labeled H and red) gene set in each mammal’s low and high recombining regions on their respective X chromosome (x-axis). Credit: Silhouettes were reproduced from PhyloPic (https://www.phylopic.org/) under a CC0 1.0 Universal Public Domain license unless otherwise stated; armadillo and pig, created by Stephen Traver; cow, created by Mozillan; dog, created by Margot Michaud; elephant, created by Ignorant Atheist; mouse, rhino, created by Christophe Mallet. Alt text: Bar graphs showcasing the percentage of autosomal (above) or X chromosome (below) genes retained in ancestral recombining regions. Each mammal has a bar graph for both high-and low-recombining regions.

## Discussion

Multiple studies have aimed to characterize the evolutionary properties of recombination rate patterns at broad and fine scales in distantly and closely related species (Jensen-Seaman et al. 2004; Dumont and Payseur 2008, 2011; Segura et al. 2013; Stevison et al. 2016; Haenel et al. 2018). A recent comparative study used machine-learning-inferred recombination rates (Adrion et al. 2020) from population genomics data and identified extremely large, conserved regions of the X chromosome that have remained conserved throughout the evolutionary history of placental mammals (Foley et al. 2026). Here we extended these analyses to identify fine-scale autosomal regions that have maintained low- and high-recombining areas in placental mammal lineages that coalesce 105 million years ago (Foley et al. 2023).

Our approach to inferring the ancestral recombination map of the placental mammal ancestor (n=23) purposely used a representative from each superordinal clade with slowly evolving karyotypes. Previous studies have attempted to reconstruct ancestral mammalian karyotypes using species regardless of karyotype evolution rates (e.g., Kim et al. 2017; Damas et al. 2022). However, large variations in evolutionary rates within the ingroup can bias ancestral-state reconstructions, and several ancestral chromosomes inferred from earlier genome sequence comparisons showed unusual associations not predicted by Zoo-FISH results (Ferguson-Smith and Trifonov 2007; Kim et al. 2017; Damas et al. 2022). Indeed, when analyzing species with slow rates of karyotypic change, the vast majority of our ancestral chromosomes reflect conserved chromosomes and syntenic associations documented in ZOO-FISH analysis (Richard et al. 2003; Ferguson-Smith and Trifonov 2007; Svartman et al. 2012) (**Fig. 1**). However, not all well-established ancestral syntenic chromosomes and human chromosome associations were recovered with DESCHRAMBLER. These findings are unusual because most of these chromosomes are intact in the species used for our reconstruction, making it unexpected that a reconstructed ancestor would lack them. However, these artifacts can be attributed to chromosomes and chromosome regions that are highly rearranged across most descendant species, where DESCHRAMBLER cannot form a contiguous ancestral block from the extensively fragmented genomic alignments (Kim et al. 2017).

The expanded repertoire of recombination maps for non-model organisms produced as part of this study represents a valuable resource for comparative genomics. Resolved maps provide crucial data needed for linkage and association studies, which are typically uncommon beyond model or domestic organisms (Palsson et al. 2025). Accurate recombination rates can also enhance population-genomic estimates of population history, mutation rates, and natural selection, which are key to informing conservation assessments of understudied species such as the sloth and aardvark (Kovacs et al. 2025). Our ReLERNN-based recombination maps exhibit the expected features of the autosomal recombination landscape, as predicted by meta-analyses of genetic maps inferred using traditional methods (Haenel et al. 2018). In longer chromosomes, high-recombination regions are maintained toward the ends, while low-recombination regions tend to be located near the center. Further, shorter chromosomes primarily consist of highly recombinant regions. Moreover, phylogenomic trees constructed from ALR regions recovered the species tree for several clades known to hybridize, underscoring the broader utility of the inferred ancestral recombination map for mammalian phylogenomics.

PhyloP scores indicative of strong purifying selection (Burbrink et al. 2025; Foley et al. 2026) characterized ALR genomic regions. Reduced recombination in these genes prevents the disruption of advantageous allelic combinations critical for essential organismal functions (Ritz et al. 2017). Previous studies of ancestral chromosome architecture have identified large homologous synteny blocks (HSBs), which are enriched for genes involved in development and the nervous system (Larkin et al. 2009; Damas et al. 2022). They concluded that these genomic blocks are selected to remain largely unchanged to preserve fundamental processes (Larkin et al. 2009; Damas et al. 2022). In support of these observations, we found that conserved, syntenic low-recombining regions were enriched for conserved pathways, including DNA repair, DNA recombination, WNT, and TGFβ signaling (**Fig. 5** and **Table S5**).

In contrast, regions involved in chromosomal rearrangements, called evolutionary breakpoint regions (EBRs), are enriched for genes involved in the sensory and immune systems (Larkin et al. 2009; Damas et al. 2022). Natural selection thus acts on adaptive and innovative configurations within these regions. Genes related to immune functions have previously been found enriched in regions of high recombination in *Myotis* bats (Foley et al. 2024), humans (Vandiedonck and Knight 2009), and chickens (Fulton et al. 2016). Interestingly, we observe several gene pathway categories enriched for processes similar to those in AHR, including inflammatory response and regulation of stimulus responses (**Fig. 5**). We hypothesize that AHR and ALR regions may be primed to facilitate adaptation, in which novel alleles are more readily brought together or deleterious mutations more effectively purged, evolving in a lineage-specific manner (Ritz et al. 2017). Interestingly, we also observed that different components of individual immune-related hallmark processes are distributed across ancestrally low and high regions. We hypothesize that movement of individual genes within a pathway to conserved high or low regions through chromosome rearrangement may represent a path through which extant species may finely adapt conserved molecular processes like immune responses. For example, Interferon pathway genes skewed towards ALR regions, whereas interleukins skewed towards AHR regions. Interferon genes evolved before interleukins, which may help explain the opposing evolutionary constraints that shaped these dynamics. In this way, our analyses have the potential to help researchers identify core-conserved components of important pathways and components of pathways more amenable to change, which may better guide human health interventions.

When we broadened our sampling of mammalian genomes with variable karyotype evolution, we found that a sizable percentage of autosomal genes from the ALR and AHR gene sets remain low and high, despite including species with a faster rate of chromosome evolution (**Fig. 6A**). However, we observed no genes that have remained in low recombining regions across all 13 mammals, whereas a small group of genes have remained high in all mammals sampled. This contradicts our expectation that natural selection would preferentially preserve regions of low recombination because of their importance in fundamental processes, such as those conserved within HSBs (Larkin et al. 2009; Damas et al. 2022). This also contrasts with the X chromosome, where large coldspots up to ∼50Mb were conserved, although they were disrupted in lineages with rearranged X chromosomes (i.e., mouse, cattle, *Myotis* bats) (**Fig. 6B**; Foley et al. 2026).

While this study focused on genomic sequence and gene content, future investigations would benefit from exploring the roles of regulatory elements and epigenetics in relation to recombination-based annotations, as these become increasingly available across many mammalian species. Extensive work has been done to understand hotspot regulation and distribution (Paigen and Petkov 2010). *Cis*- and *trans*-acting factors related to PRDM9 have been shown to direct recombination hot-spot activity in several mammalian species (Parvanov et al. 2010; Ma et al. 2015; Stevison et al. 2016). PRDM9-independent recombination hotspots have been characterized near promoters in boreoeutherian mammals with functional PRDM9 (e.g., humans, mice, cattle) and in those lacking it (e.g., canids; Joseph et al. 2024). Moreover, Joseph et al. (2024) show that hypomethylation patterns are associated with increased recombination in some mammalian species, while a deficit of recombination in others. Evaluating how these different aspects affect the conservation and diversification of recombination rates will provide a better understanding of how natural selection has shaped the recombination landscape of placental mammals.

## Materials and Methods

### Data

The list of genome assemblies used in this study, along with their NCBI accession numbers and sources, is provided in **Supplemental Table S7**. We reconstructed the genomes of the aardvark and the sloth because they lacked high-quality, chromosomal-level assemblies. PacBio SMRT libraries were size-selected (> 20 kb) from high molecular weight DNA and sequenced on the PacBio Sequel II to 59x for the aardvark and 68x for the sloth. Illumina libraries were size-selected (300 bp) and run on the Illumina NovaSeq 600, with a coverage of 132x for the aardvark and 82x for the sloth. PacBio reads were filtered with a 7-kb cutoff in Aardvark and a 15-kb cutoff in Sloth.

Reads were assembled into contigs using Flye v2.8.3-b1695 for the aardvark (Kolmogorov et al. 2019) and NextDenovo v2.2-beta.0 for the sloth (Hu et al. 2024). Aardvark and sloth contigs were scaffolded into a chromosomal-level assembly using Hi-C sequencing (Dudchenko et al. 2017). Hi-C sequencing data were obtained from the DNA Zoo Consortium (PRJNA512907; SRR8616860 for aardvark and SRR8616918 for sloth). A map of Hi-C contacts for downstream scaffolding was generated using the Juicer v1.6 algorithm with the restriction enzyme MboI (Durand et al. 2016b). The 3D-DNA v180419 pipeline scaffolded the contigs and corrected any misassemblies using the Hi-C contact map (Dudchenko et al. 2017). Misassemblies were manually inspected and corrected using Juicebox Assembly Tools v1.11.08 (Dudchenko et al. 2018; Durand et al. 2016a). Published chromosome painting data were used to identify chromosomes (aardvark: Yang et al. 2003; sloth: Svartman et al. 2006). This was done by aligning the aardvark and sloth assemblies to the human assembly (T2T-CHM13v2.0) using nucmer with default settings (mummer v4.0.0rc1; Marçais et al. 2018). Dot plots were visualized on the interactive dot plot viewer, Dot (https://github.com/MariaNattestad/dot). Adaptor sequences were identified and removed using FCS-adaptor v0.2.3 (Astashyn et al. 2024).

Initial long-read assemblies were polished with Illumina reads. For the aardvark, we used ntHits v0.1.0 (Mohamadi et al. 2020) and ntedit v1.3.5 (Warren et al. 2019). For the sloth, we ran two rounds of Hapo-G v1.3.4 (Aury et al. 2021), using a BAM file generated with BWA v0.7.17 (Li 2013) and SAMtools v1.13 (Danecek et al. 2021). A k-mer database was first built using Meryl v1.3 (Rhie et al. 2020) to obtain a quality value score (QV) from Illumina reads for both species. QV was calculated for each assembly using its k-mer database and Merqury v1.3 (Rhie et al. 2020). Genome completeness was evaluated using Compleasm v0.2.06 with the mammalia_odb10 library (Huang and Li 2023). In addition, QUAST v5.3.0 was used to assess the quality of the genome assemblies (Gurevich et al. 2013). Gene annotations for both assemblies were generated using Liftoff v1.6.3 (Shumate and Salzberg 2020) to lift over gene annotations from the OryAfe1.0 aardvark assembly and from the closely related sister taxa of the sloths, Linnaeus’s two-toed sloth (*Choloepus didactylus*; mChoDid1.pri).

Lastly, we assembled a mitochondrial genome assembly for the aardvark. The mitochondrial genome was assembled by using minimap2 v2.23 (Li 2018) to map the long-read data against the mitochondrial sequence from a previously published aardvark assembly (OryAfe1.0). Samtools v1.12 (Danecek et al. 2021) created a BAM file from the SAM file outputted by the aligner. A consensus sequence was generated using Geneious Prime 2022.1.1 (https://www.geneious.com). The mitochondrial sequence in our assembly was identified and assembled by blasting (BLAST+ v2.12.0) our consensus sequence against our assembly (Camacho et al. 2009).

### Synteny Analysis

Synteny plots were generated using two different approaches from the R package, GENESPACE (Lovell et al. 2022). First, gene-based syntenic blocks were identified using gene annotations (gff) and a peptide fasta file (translated cds) using GENESPACE v1.3.1. Second, sequence-based syntenic blocks were identified using whole-genome alignments with GENESPACEv1.4.1. GENESPACE was run on our five mammals used in the ancestral reconstruction (i.e., human, domestic cat, aardvark, Hoffmann’s two-toed sloth, and the blue whale) and the thirteen mammal-based reconstruction of ancestral recombination rates. We had to lift annotations for the nine-banded armadillo and the white rhino for the thirteenth mammal comparison, as they lacked gene annotations (Shumate and Salzberg 2020). For the nine-banded armadillo, we lifted annotations from the mDasNov1.hap2 nine-banded armadillo assembly. For the white rhino, we lifted annotations from the CerSimSim1.0 white rhino assembly. AGAT v0.9.2 (Dainat 2022) converted genes from gene annotation files into amino acid sequences for the aardvark, the sloth, the nine-banded armadillo, and the white rhino. GENESPACE v1.3.1 was run with default parameters using the following programs: MCScanX v2022.10.31 (Wang et al. 2012), DIAMOND v2.0.15 (Buchfink et al. 2021), and OrthoFinder 2.5.4 (Emms and Kelly 2019). Under v1.4.1, fasta-formatted genome assemblies were restricted to chromosome-level scaffolds for pairwise GENESPACE comparisons. GENESPACE v1.4.1 ran with the following programs: MCScanX v2022.10.31 (Wang et al. 2012), DIAMOND v.2.1.0 (Buchfink et al. 2021), and minimap2 v2.24 (Li 2018).

### Ancestor Reconstruction

The following genomes were used in the reconstruction of the ancestral placental mammal genome: human (*Homo sapiens*, GRCh38.p14), domestic cat (*Felis catus*, Fca126_mat1.0), blue whale (*Balaenoptera musculus*, mBalMus1.pri.v3), aardvark (*Orycteropus afer*, this study), and Hoffmann’s two-toed sloth (*Choloepus hoffmanni*, this study). We used the DESCHRAMBLER algorithm to reconstruct the ancestral karyotype of placental mammals (Kim et al. 2017). Due to its high accuracy and completeness, we used the human genome as the reference to build the reconstruction. As the closest sister taxon to placental mammals, marsupials are not an ideal karyotype for ancestral state reconstruction, given the more than 160 million years of independent divergence from placental mammals (Phillips et al. 2009; Luo et al. 2011). Furthermore, marsupial karyotypes are biased towards lower chromosome numbers, which could lower ancestral chromosome numbers (Deakin et al. 2012). To avoid this bias, the aardvark was used as the outgroup (Kim et al. 2017; Damas et al. 2022).

We generated pairwise alignments between the human genome (the reference) and the other four mammals using Lastz_32 v1.04.15 (Harris 2007). Before aligning the genomes, we removed unplaced scaffolds, mitochondrial sequences, and Y chromosome sequences. After generating the alignments, we converted them into chain and net files using GenomeAlignmentTools v2019.11.20 with the following commands in order: axtChain, chainSplit, chainSort, chainPreNet, and chainNet (Kent et al. 2003). To run DESCHRAMBLER, the required inputs are chain and net files, a phylogenetic tree, a configuration, and a parameter file. The tree (((Human:0.1806,(Whale:0.1246,Domestic_Cat:0.1709):0.0435):0.0186,Hoffmann_Two_Toed_ Sloth:0.175)@:0.0186332,Aardvark:0.2145); was modified from Foley et al. (2023). Block resolution and minimum adjacency score were set at 500,000 base pairs (bp) and 0.0001, respectively. In our sensitivity analysis, we varied the block resolution (300,000 or 500,000 bp), the reference (human or sloth), and the outgroup (aardvark, sloth, or a combination of aardvark and sloth). The ancestral karyotype was illustrated using the R package, syntenyPlotteR (Quigley et al. 2023).

### Generation of recombination maps

In addition to previously published recombination maps (Foley et al 2026), additional maps were generated for the sloth and aardvark. Raw Illumina reads for the sloth and aardvark were filtered using Trim Galore! v0.6.5 (Krueger et al. 2015) and mapped to their respective reference genomes using bwa-mem v0.7.17 (Li et al. 2009; Li et al. 2013) using the -M and -R parameters. Mapping results were summarized using the Qualimap function bamqc (García-Alcalde et al. 2012). Samtools v1.9 (Li et al. 2009) was used to remove duplicate reads. Local realignment and variant calling were performed using GATK v4.1.2 (McKenna et al. 2010; DePristo et al. 2011; Poplin et al. 2017; Van der Auwera et al. 2013). Variants were called, and all samples were jointly genotyped. Repetitive sequences in each genome were identified using RepeatMasker v4.3.2 (Smit et al. 2013-2015) and the Zoonomia repeat library (Osmanski et al. 2023). Variants in repeatmasked regions were masked using GATK. Variants were further filtered, removing variants within 5bps of an indel and those which did not meet the following quality criteria -e’%QUAL<30 | INFO/DP<16 | INFO/DP>62 | QD<2 | FS>60 | SOR>10 | ReadPosRankSum <-8 | MQRankSum <-12.5 | MQ<40’ in bcftools (Danecek and McCarthy 2017). VCFtools were used to remove indels (Danecek et al. 2011). ReLERNN, a deep learning approach that uses recurrent neural networks, was used to model the genome-wide recombination rate for each species. Given that empirical mutation rates for the aardvark and sloth are unknown, the average mammalian mutation rate of 2.2e-9 was used in the analyses (Kumar & Subramanian 2002; Foley et al. 2026). ReLERNN was run using the simulate, train, predict, and bscorrect modules with default settings. Finally, inferred recombination rates were averaged into 2 Mb blocks with 50kb sliding windows for use in downstream analyses. The sex-averaged human recombination map was taken from Halldorsson et al. (2019). Due to the low sequence divergence of the recombination desert in the middle of the X chromosome, there was a lack of recombination rate data for some mammals. Given that this region likely has low recombination rates, we set the recombination maps of the X chromosome in the domestic cat, Asian elephant, pig, nine-banded armadillo, and cow to 0 in data-free windows within the recombination desert.

### Identifying regions that retained a low or high recombination rate

The four mammals used in this analysis were the human, domestic cat, aardvark, and Hoffmann’s two-toed sloth. The analysis was restricted to the autosomes and X chromosomes and excluded the Y chromosome, mitochondria, or any unplaced scaffolds. First, BEDTools v2.30.0 generated 1.5 Mbp windows for each mammalian chromosome based on its size (Quinlan and Hall 2010). Windows from each genome were combined, and for each species recombination map, BEDTools was used to map averaged recombination rates for each window (Quinlan and Hall 2010). Each species’ average window rate was ranked from highest to lowest and divided into four rate categories (quartiles). Using a combination of custom R scripts and manual curation, we identified syntenic regions supported by our ancestral placental mammal reconstruction, GENESPACE results, and nucmer alignments that have retained either low or high recombination rates.

To identify these syntenic regions, we first narrowed them down based on our ancestral placental mammal reconstruction and GENESPACE results. These regions are the chromosomes that have remained stable across millions of years of mammalian evolution. To further narrow the regions that have retained low or high recombination rates, we examined the collinearity of human- and cat-based windows within the lowest- or highest-recombination-rate categories relative to the rest of mammals. Nucmer and GENESPACE v1.4.1 alignments were used to detect collinearity among the different genomes. We then identified the recombination-rate categories for each collinear region. A region was classified as low or high if it contained either a recombination extreme, the next rate category above, or a mixture of the two. If a single region contained both low- and high-rate categories, the category present in 50% or more of the collinear region was assigned. Once rate categories were assigned, a region was identified as retaining a low or high recombination rate if four of the mammals, or three of the four mammals, possessed the same category. Per-rate coordinates were compiled using the human and cat window references (**Table S2**).

For the final coordinates, two regions were merged if they were within a megabase of each other, and coordinates were edited to prevent overlap between ALR and AHR coordinates (e.g., a cutoff in the middle of the overlap to denote ALR and AHR). The sloth X chromosome regions were excluded from the final analysis due to the complexities of their sex chromosomes (Corin-Frederic 1969; Jorge et al. 1985). Recombination maps displaying extant (**Fig. S1-S4**) or ancestral (**Fig. 3** and **Fig. S5-S8**) recombination rates were generated using the R package karyoploteR (Gel and Serra 2017). To generate **Fig. 3**, we had to map the ancestral recombination landscape to our reconstructed ancestral karyotype. We used human-conserved recombination coordinates to map the human syntenic regions that comprise the ancestral placental mammal karyotype. ChatGPT vGPT-4o was prompted to generate the following formula:

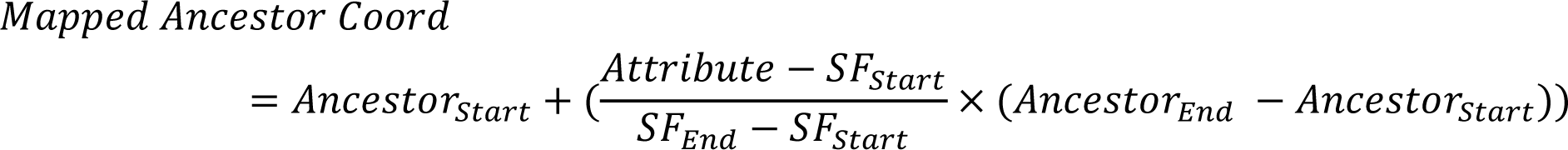

Attribute refers to the start or end coordinates of the conserved recombination rate region. SF is the syntenic fragment of the human within the reconstructed ancestor. An ancestor refers to the coordinates of the reconstructed ancestral placental mammal chromosomes.

### Gene set enrichment analysis

We performed a gene set enrichment analysis to test whether genes within our conserved regions, using the Web-based Gene Set Analysis Toolkit (WebGestalt) and STRING, were enriched for functional categories (Szklarczyk et al. 2023; Elizarraras et al. 2024). We performed two analyses: with genes from ALR and AHR. To avoid reference bias, we utilized genes shared between humans and cats. Custom R scripts extracted genes found within those regions in human and cat, and the gene list was manually edited to remove any genes that were not one-to-one orthologs between the two species.

For our WebGestalt analysis, we performed an overrepresentation analysis using the Gene Ontology (Biological Process, Cellular Component, and Molecular Function) and pathway databases (KEGG, Panther, Reactome, and Wikipathways). We applied the Benjamini-Hochberg (BH) false discovery rate correction with a significance threshold of p < 0.05. To reduce redundancy, we chose a weighted set cover (**Fig. 5A**). For our STRING analysis, we submitted the list of genes to STRING’s multiple protein search GUI (https://string-db.org/) with human as our organism of interest (Szklarczyk et al. 2023). The enrichment display was chosen with default settings, including a maximum FDR of <= 0.05. To reduce redundancy, we merge rows based on term similarity >= 0.5 (**Fig. 5A**).

### Comparison of recombination rates across mammals

We compared the recombination landscape of 13 divergent mammals with different karyotype evolutionary histories with the regions we identified as having low or high recombination rates. We utilized each mammal gene set with recombination rate information for the comparison. The thirteen mammals used in this analysis were the human (*Homo sapiens*, GRCh38.p14), domestic cat (*Felis catus*, F.catus_Fca126_mat1.0), aardvark (*Orycteropus afer*, TAMU_Oaf_1.0), Hoffmann’s two-toed sloth (*Choloepus hoffmanni*, TAMU_Cho_1.0), blue whale (*Balaenoptera musculus*, mBalMus1.pri.v3), cow (*Bos taurus*, ARS-UCD2.0), nine-banded armadillo (*Dasypus novemcinctus*, mDasNov1.1.hap2), greater mouse-eared bat (*Myotis myotis*, mMyoMyo1.p), pig (*Sus scrofa*, Sscrofa11.1), Asian elephant (*Elephas maximus*, mEleMax1), the dog (*Canis lupus familiaris*, Dog10K_Boxer_Tasha), house mouse (*Mus musculus*, GRCm39) and the white rhino (*Ceratotherium simum*, ASM2365373v1).

First, we wanted to resolve as many unnamed genome annotations, or LOCs, as possible to increase the number of named genes available for analysis. LOCs from genomes of interest were assigned to orthogroups using OrthoFinder v2.5.4 (Emms and Kelly 2019). Using custom R scripts and manual curation, LOCs were identified by gene symbol if the same gene was found to be an ortholog in humans, dogs, mice, or at least two mammals. In the case of dog and mouse, LOCs were identified when an ortholog was found in both species (human and mouse, or human and dog).

Next, genes were assigned to recombination rate categories. BEDtools v2.30.0 divided and averaged each mammalian recombination map into 1.5 Mb windows, following the procedure described above (Quinlan and Hall 2010). Custom R scripts were used to rank and assign windows to recombination quartiles. In some cases, we identified genes that fell into two different rate categories. This was resolved by their random assignment to one of the two rate categories.

To calculate the percentage of genes retained in the ancestral recombining regions, we first compiled a list of genes shared by all mammals in our study. This was done to avoid biased percentages arising from missing data, whether a gene was absent in a particular species or not annotated. The number of genes for autosomal low, autosomal high, X low, and X high were 1,138, 1,565, 76, and 17, respectively. We calculated the number of genes retained in the low-and high-recombining regions of the autosomes and the X chromosome using a custom R script and the gene databases generated above. We output the retained genes for each mammal to identify which genes remained in all 13 species’ low- and high-recombining genomic regions (**Fig. 6**; **Table S5**).

### Phylogenomics of ancestrally high- and low-recombining regions in mammals

To examine the phylogenomic properties of the ALR and AHR regions, we obtained human (hg38)-referenced coordinates for ancestrally conserved high- and low-recombining regions (see **Table S2** under “Final Human Coords”). Fasta alignments corresponding to these coordinates were extracted from the human-referenced Zoonomia alignment, which comprises 240 placental mammal species. Alignments were filtered to remove any alignment column with greater than 10% missingness using trimAl v1.4.1 (Capella-Gutierrez et al. 2009). Additionally, poorly aligned regions were further filtered using a sliding-window alignment filtering script (https://github.com/VCMason/Foley2021) with the parameters described in Foley et al. 2023. Alignments were then concatenated to form 1) an ancestrally low and 2) an ancestrally high recombining dataset for further analysis. Because the resulting datasets were too large to efficiently compute a maximum-likelihood tree, we randomly selected 100,000 alignment columns from each dataset for further analysis. Maximum Likelihood (ML) trees were generated using IQ-TREE2 v2.2.1 (Minh et al. 2020) under a GTR model of sequence evolution and the GHOST model of rate variation (GTR*H4). The highest-scoring ML tree for both datasets was evaluated with 1000 bootstrap replicates using the ultrafast bootstrap approximation.

Trees derived from ALR and AHR regions were manually inspected to identify topological differences between trees from these regions. To determine if there were systematic differences in branch lengths between conserved high and low regions, branch lengths from both datasets were extracted using ape v5.8.1 in R and plotted (Paradis and Schliep 2019). Similarly, phytools v2.5.1 was used to extract all node heights found in common between the two topologies, and their differences were investigated (Revell 2011).

Publicly available PhyloP scores from the human-referenced Zoonomia alignment were downloaded from https://zoonomiaproject.org/the-data/. In general, positive PhyloP scores indicate conservation, zero values represent neutrally evolving sites, and negative numbers indicate accelerated sites (Pollard et al. 2010; Hubisz et al. 2011). PhyloP scores corresponding to the human (hg38) reference coordinates for ancestrally conserved high- and low-recombining regions were collected using bedtools v2.30.0 intersect (Quinlan and Hall 2010) and plotted.

### Hallmark Gene Set Collection

Given that PhyloP scores were more conserved in ancestrally low recombining regions, we sought to quantify the relative distribution of genes across key biological pathways relative to ancestrally conserved recombination regions. In this way, we hope to identify genes that may represent “core conserved” components of a pathway (ancestrally low recombining) and those genes freer to vary (ancestrally high recombining). To do this, we queried the gene sets corresponding to conserved high and low regions on autosomes (**Table S5**) against the Hallmark Gene Set Collection (Liberzon et al 2015). These 50 gene sets represent well-defined biological states known to exhibit coherent expression within pathways, e.g., inflammation, spermatogenesis, DNA repair, oxidative phosphorylation, and angiogenesis. To determine whether genes in a given pathway were significantly skewed toward ancestrally low- or high-recombining regions, we compared the ratios of genes in each pathway across regions and controlled for the number of genes in each pathway. A chi-squared test was used to determine whether genes in a pathway were significantly skewed toward ancestrally high or low regions, and multiple corrections were accounted for using the Benjamini-Hochberg approach in R.

## Acknowledgments

We thank John Lovell for providing the pre-released version of GENESPACE v1.4.1 for this analysis. We acknowledge the DOE Joint Genome Institute, which funds the development of GENESPACE. The tissue sample from the muscle of a Hoffmann’s two-toed sloth was provided by Angelo State Natural History Collections (Loren Ammerman). The aardvark genome was produced with DNA provided by Dr. Terrence Robinson (Meredith et al. 2011). We thank the following students for their help with some analyses for this study during the Fall 2024 Aggie Research Project (ARP): Juiyun Tsai, Saveer Ramnani, Madi Shojaei, and Charne Du Plooy. Hi-C sequencing data for the aardvark and Hoffmann’s two-toed sloth are used with permission from the DNA Zoo Consortium (dnazoo.org). Portions of this research were conducted using the advanced computing resources provided by Texas A&M High Performance Research Computing.

## Data Availability

The Hoffmann’s two-toed sloth and aardvark BioProject are accessible at the NCBI BioProject (BioProject; https://www.ncbi.nlm.nih.gov/bioproject/) under the accession number PRJNA1073880. Whole-genome Illumina and PacBio sequencing reads of the aardvark and sloth have been submitted to the NCBI Sequence Read Archive (SRA; https://www.ncbi.nlm.nih.gov/sra/) under accession numbers SRR27888359, SRR27888358, SRR27888357, and SRR27888356. The aardvark and Hoffmann’s two-toed sloth’s genome assemblies have been submitted to the NCBI GenBank under biosample accession numbers SAMN39842544 and SAMN39842543, respectively.

Custom R scripts used in the analysis of this study are available at GitHub (https://github.com/ichilde/Recombination-Rate-Manuscript/).

## Author Contributions

IRC: Data curation; Formal analysis; Investigation; Methodology; Software; Validation; Visualization; Writing—original draft; Writing—review & editing. NMF: Data curation; Formal analysis; Investigation; Methodology; Resources; Validation; Visualization; Writing—review & editing. KRB: Formal analysis; Investigation; Data curation. WJM: Conceptualization; Funding acquisition; Investigation; Methodology; Project administration; Resources; Supervision; Writing—review & editing.

## References

1. Adrion JR, Galloway JG, Kern AD. 2020. Predicting the Landscape of Recombination Using Deep Learning. Mol Biol Evol 37: 1790–1808. 10.1093/molbev/msaa038.

2. Astashyn A, Tvedte ES, Sweeney D, Sapojnikov V, Bouk N, Joukov V, Mozes E, Strope PK, Sylla PM, Wagner L, et al. 2024. Rapid and sensitive detection of genome contamination at scale with FCS-GX. Genome Biol 25: 60. 10.1186/s13059-024-03198-7.

3. Aury J-M, Istace B. 2021. Hapo-G, haplotype-aware polishing of genome assemblies with accurate reads. NAR Genom Bioinform 3: lqab034. 10.1093/nargab/lqab034.

4. Axelsson E, Webster MT, Ratnakumar A, LUPA Consortium, Ponting CP, Lindblad-Toh K. 2012. Death of PRDM9 coincides with stabilization of the recombination landscape in the dog genome. Genome Res 22: 51–63. 10.1101/gr.124123.111.

5. Baudat F, Buard J, Grey C, Fledel-Alon A, Ober C, Przeworski M, Coop G, de Massy B. 2010. PRDM9 is a major determinant of meiotic recombination hotspots in humans and mice. Science 327: 836–840. 10.1126/science.1183439.

6. Baudat F, Imai Y, de Massy B. 2013. Meiotic recombination in mammals: localization and regulation. Nat Rev Genet 14: 794–806. 10.1038/nrg3573.

7. Brashear WA, Bredemeyer KR, Murphy WJ. 2021. Genomic architecture constrained placental mammal X Chromosome evolution. Genome Res 31: 1353–1365. 10.1101/gr.275274.121.

8. Buchfink B, Reuter K, Drost H-G. 2021. Sensitive protein alignments at tree-of-life scale using DIAMOND. Nat Methods 18: 366–368. 10.1038/s41592-021-01101-x.

9. Burbrink FT, DeBaun D, Foley NM, Murphy WJ. 2025. Recombination-aware phylogenomics. Trends Ecol Evol 0. 10.1016/j.tree.2025.06.011.

10. Camacho C, Coulouris G, Avagyan V, Ma N, Papadopoulos J, Bealer K, Madden TL. 2009. BLAST+: architecture and applications. BMC Bioinformatics 10: 421. 10.1186/1471-2105-10-421.

11. Capella-Gutierrez S, Silla-Martinez JM, Gabaldon T. 2009. trimAl: a tool for automated alignment trimming in large-scale phylogenetic analyses. Bioinformatics 25: 1972–1973. 10.1093/bioinformatics/btp348.

12. Chakravarti A, Elbein SC, Permutt MA. 1986. Evidence for increased recombination near the human insulin gene: implication for disease association studies. Proc Natl Acad Sci U S A 83: 1045–1049. 10.1073/pnas.83.4.1045.

13. Christmas MJ, Kaplow IM, Genereux DP, Dong MX, Hughes GM, Li X, Sullivan PF, Hindle AG, Andrews G, Armstrong JC, et al. 2023. Evolutionary constraint and innovation across hundreds of placental mammals. Science 380: eabn3943. 10.1126/science.abn3943.

14. Corin-Frederic J. 1969. Les formules gonosomiques dites aberrantes chez les Mammifères Euthériens. Chromosoma 27: 268–287. 10.1007/BF00326165.

15. Dainat J. 2022. Another Gtf/Gff Analysis Toolkit (AGAT): Resolve interoperability issues and accomplish more with your annotations. Plant and Animal Genome XXIX Conference. https://github.com/NBISweden/AGAT.

16. Damas J, Corbo M, Kim J, Turner-Maier J, Farré M, Larkin DM, Ryder OA, Steiner C, Houck ML, Hall S, et al. 2022. Evolution of the ancestral mammalian karyotype and syntenic regions. Proc Natl Acad Sci U S A 119: e2209139119. 10.1073/pnas.2209139119.

17. Danecek P, Auton A, Abecasis G, Albers CA, Banks E, DePristo MA, Handsaker RE, Lunter G, Marth GT, Sherry ST, et al. 2011. The variant call format and VCFtools. Bioinformatics 27: 2156–2158. 10.1093/bioinformatics/btr330.

18. Danecek P, Bonfield JK, Liddle J, Marshall J, Ohan V, Pollard MO, Whitwham A, Keane T, McCarthy SA, Davies RM, et al. 2021. Twelve years of SAMtools and BCFtools. Gigascience 10. 10.1093/gigascience/giab008.

19. Danecek P, McCarthy SA. 2017. BCFtools/csq: haplotype-aware variant consequences. Bioinformatics 33: 2037–2039. 10.1093/bioinformatics/btx100.

20. Deakin JE, Graves JAM, Rens W. 2012. The evolution of marsupial and monotreme chromosomes. Cytogenet Genome Res 137: 113–129. 10.1159/000339433.

21. Delgado CLR, Waters PD, Gilbert C, Robinson TJ, Graves JAM. 2009. Physical mapping of the elephant X chromosome: conservation of gene order over 105 million years. Chromosome Res 17: 917–926. 10.1007/s10577-009-9079-1.

22. DePristo MA, Banks E, Poplin R, Garimella KV, Maguire JR, Hartl C, Philippakis AA, del Angel G, Rivas MA, Hanna M, et al. 2011. A framework for variation discovery and genotyping using next-generation DNA sequencing data. Nat Genet 43: 491–498. 10.1038/ng.806.

23. Dudchenko O, Batra SS, Omer AD, Nyquist SK, Hoeger M, Durand NC, Shamim MS, Machol I, Lander ES, Aiden AP, et al. 2017. De novo assembly of the Aedes aegypti genome using Hi-C yields chromosome-length scaffolds. Science 356: 92–95. 10.1126/science.aal3327.

24. Dudchenko O, Shamim MS, Batra SS, Durand NC, Musial NT, Mostofa R, Pham M, St Hilaire BG, Yao W, Stamenova E, et al. 2018. The Juicebox Assembly Tools module facilitates de novo assembly of mammalian genomes with chromosome-length scaffolds for under $1000. *bioRxiv* 254797. https://www.biorxiv.org/content/10.1101/254797.

25. Dumont BL, Payseur BA. 2008. Evolution of the genomic rate of recombination in mammals. Evolution 62: 276–294. 10.1111/j.1558-5646.2007.00278.x.

26. Dumont BL, Payseur BA. 2011. Evolution of the genomic recombination rate in murid rodents. Genetics 187: 643–657. 10.1534/genetics.110.123851.

27. Dunnum JL, Salazar-Bravo J, Yates TL. 2001. The Bolivian bamboo rat, Dactylomys boliviensis (Rodentia: Echimyidae), a new record for chromosome number in a mammal. Acta Histochemica 103: 121–126. https://scholar.google.com/citations?user=7O5fxgYAAAAJ&hl=en&oi=sra.

28. Durand NC, Robinson JT, Shamim MS, Machol I, Mesirov JP, Lander ES, Aiden EL. 2016a. Juicebox Provides a Visualization System for Hi-C Contact Maps with Unlimited Zoom. Cell Syst 3: 99–101. 10.1016/j.cels.2015.07.012.

29. Durand NC, Shamim MS, Machol I, Rao SSP, Huntley MH, Lander ES, Aiden EL. 2016b. Juicer Provides a One-Click System for Analyzing Loop-Resolution Hi-C Experiments. Cell Syst 3: 95– 98. 10.1016/j.cels.2016.07.002.

30. Elizarraras JM, Liao Y, Shi Z, Zhu Q, Pico AR, Zhang B. 2024. WebGestalt 2024: faster gene set analysis and new support for metabolomics and multi-omics. Nucleic Acids Res 52: W415–W421. https://academic.oup.com/nar/article-pdf/52/W1/W415/58435853/gkae456.pdf.

31. Emms DM, Kelly S. 2019. OrthoFinder: phylogenetic orthology inference for comparative genomics. Genome Biol 20: 238. 10.1186/s13059-019-1832-y.

32. Ferguson-Smith MA, Trifonov V. 2007. Mammalian karyotype evolution. Nat Rev Genet 8: 950–962. 10.1038/nrg2199.

33. Foley NM, Rasulis RG, Wani Z, Mendoza Cerna MN, Figueiró HV, Koepfli KP, Raudsepp T, Murphy WJ. 2026. An ancient recombination desert is a speciation supergene in placental mammals. Nature 649: 1228–1236. 10.1038/s41586-025-09740-2.

34. Foley NM, Mason VC, Harris AJ, Bredemeyer KR, Damas J, Lewin HA, Eizirik E, Gatesy J, Karlsson EK, Lindblad-Toh K, et al. 2023. A genomic timescale for placental mammal evolution. Science 380: eabl8189. 10.1126/science.abl8189.

35. Foley NM, Harris AJ, Bredemeyer KR, Ruedi M, Puechmaille SJ, Teeling EC, Criscitiello MF, Murphy WJ. 2024. Karyotypic stasis and swarming influenced the evolution of viral tolerance in a species-rich bat radiation. Cell Genom 4: 100482. http://www.cell.com/article/S2666979X23003348/abstract.

36. Fulton JE, McCarron AM, Lund AR, Pinegar KN, Wolc A, Chazara O, Bed’Hom B, Berres M, Miller MM. 2016. A high-density SNP panel reveals extensive diversity, frequent recombination and multiple recombination hotspots within the chicken major histocompatibility complex B region between BG2 and CD1A1. Genet Sel Evol 48: 1. 10.1186/s12711-015-0181-x.

37. García-Alcalde F, Okonechnikov K, Carbonell J, Cruz LM, Götz S, Tarazona S, Dopazo J, Meyer TF, Conesa A. 2012. Qualimap: evaluating next-generation sequencing alignment data. Bioinformatics 28: 2678–2679. 10.1093/bioinformatics/bts503.

38. Gel B, Serra E. 2017. karyoploteR: an R/Bioconductor package to plot customizable genomes displaying arbitrary data. Bioinformatics 33: 3088–3090. 10.1093/bioinformatics/btx346.

39. Graphodatsky AS, Trifonov VA, Stanyon R. 2011. The genome diversity and karyotype evolution of mammals. Mol Cytogenet 4: 22. 10.1186/1755-8166-4-22.

40. Gurevich A, Saveliev V, Vyahhi N, Tesler G. 2013. QUAST: quality assessment tool for genome assemblies. Bioinformatics 29: 1072–1075. 10.1093/bioinformatics/btt086.

41. Haenel Q, Laurentino TG, Roesti M, Berner D. 2018. Meta-analysis of chromosome-scale crossover rate variation in eukaryotes and its significance to evolutionary genomics. Mol Ecol 27: 2477–2497. 10.1111/mec.14699.

42. Halldorsson BV, Palsson G, Stefansson OA, Jonsson H, Hardarson MT, Eggertsson HP, Gunnarsson B, Oddsson A, Halldorsson GH, Zink F, et al. 2019. Characterizing mutagenic effects of recombination through a sequence-level genetic map. Science 363. 10.1126/science.aau1043.

43. Harris RS. 2007. Improved pairwise alignment of genomic DNA. PhD, The Pennsylvania State University, Centre County, Pennsylvania http://proxy.library.tamu.edu/login?url= https://www.proquest.com/dissertations-theses/improved-pairwise-alignment-genomic-dna/docview/304835295/se-2.

44. Hu J, Wang Z, Sun Z, Hu B, Ayoola AO, Liang F, Li J, Sandoval JR, Cooper DN, Ye K, et al. 2024. NextDenovo: an efficient error correction and accurate assembly tool for noisy long reads. Genome Biol 25: 107. 10.1186/s13059-024-03252-4.

45. Huang N, Li H. 2023. compleasm: a faster and more accurate reimplementation of BUSCO. Bioinformatics 39. 10.1093/bioinformatics/btad595.

46. Hubisz MJ, Pollard KS, Siepel A. 2011. PHAST and RPHAST: phylogenetic analysis with space/time models. Brief Bioinform 12: 41–51. 10.1093/bib/bbq072.

47. Jensen-Seaman MI, Furey TS, Payseur BA, Lu Y, Roskin KM, Chen C-F, Thomas MA, Haussler D, Jacob HJ. 2004. Comparative recombination rates in the rat, mouse, and human genomes. Genome Res 14: 528–538. 10.1101/gr.1970304.

48. Jorge W, Orsi-Souza AT, Best R. 1985. The somatic chromosome of Xenarthra. In The Evolution and Ecology of Armadillos, Sloths and Vermilinguas, pp. 121–129, Smithsonian Institution Press, Washington.

49. Joseph J, Prentout D, Laverré A, Tricou T, Duret L. 2024. High prevalence of PRDM9-independent recombination hotspots in placental mammals. Proc Natl Acad Sci U S A 121: e2401973121. https://www.pnas.org/doi/abs/10.1073/pnas.2401973121.

50. Kent WJ, Baertsch R, Hinrichs A, Miller W, Haussler D. 2003. Evolution’s cauldron: duplication, deletion, and rearrangement in the mouse and human genomes. Proc Natl Acad Sci U S A 100: 11484–11489. https://www.pnas.org/doi/abs/10.1073/pnas.1932072100.

51. Kim J, Farré M, Auvil L, Capitanu B, Larkin DM, Ma J, Lewin HA. 2017. Reconstruction and evolutionary history of eutherian chromosomes. Proc Natl Acad Sci U S A 114: E5379–E5388. 10.1073/pnas.1702012114.

52. Kolmogorov M, Yuan J, Lin Y, Pevzner PA. 2019. Assembly of long, error-prone reads using repeat graphs. Nat Biotechnol 37: 540–546. 10.1038/s41587-019-0072-8.

53. Kovacs TGL, Foley NM, Silver LW, McLennan EA, Hogg CJ, Ho SYW. 2025. Mutation rate estimate and population genomic analysis reveals decline of koalas prior to human arrival. *bioRxiv* 2025.05.15.654135. https://www.biorxiv.org/content/10.1101/2025.05.15.654135v1.abstract.

54. Krueger, F. Trim Galore!: A wrapper around Cutadapt and FastQC to consistently apply adapter and quality trimming to FastQ files, with extra functionality for RRBS data (Babraham Institute).

55. Kumar S, Subramanian S. 2002. Mutation rates in mammalian genomes. Proc Natl Acad Sci U S A 99: 803–808. 10.1073/pnas.022629899.

56. Lan L, Zhang X, Xie J, Lin X, Hong X, Nie W, Wang J, Su W, Yang F, He G, et al. 2026. Comparative genomics provide insights into karyotype evolution in vespertilionid bats (Vespertilionidae, Chiroptera). Mol Ecol Resour 26: e70129. 10.1111/1755-0998.70129.

57. Larkin DM, Pape G, Donthu R, Auvil L, Welge M, Lewin HA. 2009. Breakpoint regions and homologous synteny blocks in chromosomes have different evolutionary histories. Genome Res 19: 770–777. 10.1101/gr.086546.108.

58. Li G, Figueiró HV, Eizirik E, Murphy WJ. 2019. Recombination-Aware Phylogenomics Reveals the Structured Genomic Landscape of Hybridizing Cat Species. Mol Biol Evol 36: 2111–2126. 10.1093/molbev/msz139.

59. Li H. 2013. Aligning sequence reads, clone sequences and assembly contigs with BWA-MEM. *arXiv [q-bioGN]*. http://arxiv.org/abs/1303.3997.

60. Li H. 2018. Minimap2: pairwise alignment for nucleotide sequences. Bioinformatics 34: 3094–3100. 10.1093/bioinformatics/bty191.

61. Li H, Durbin R. 2009. Fast and accurate short read alignment with Burrows-Wheeler transform. Bioinformatics 25: 1754–1760. 10.1093/bioinformatics/btp324.

62. Liberzon A, Birger C, Thorvaldsdóttir H, Ghandi M, Mesirov JP, Tamayo P. 2015. The Molecular Signatures Database (MSigDB) hallmark gene set collection. Cell Syst 1: 417–425. 10.1016/j.cels.2015.12.004.

63. Lovell JT, Sreedasyam A, Schranz ME, Wilson M, Carlson JW, Harkess A, Emms D, Goodstein DM, Schmutz J. 2022. GENESPACE tracks regions of interest and gene copy number variation across multiple genomes. Elife 11. 10.7554/eLife.78526.

64. Luo Z-X, Yuan C-X, Meng Q-J, Ji Q. 2011. A Jurassic eutherian mammal and divergence of marsupials and placentals. Nature 476: 442–445. 10.1038/nature10291.

65. Ma L, O’Connell JR, VanRaden PM, Shen B, Padhi A, Sun C, Bickhart DM, Cole JB, Null DJ, Liu GE, et al. 2015. Cattle sex-specific recombination and genetic control from a large pedigree analysis. PLoS Genet 11: e1005387. 10.1371/journal.pgen.1005387.

66. Marçais G, Delcher AL, Phillippy AM, Coston R, Salzberg SL, Zimin A. 2018. MUMmer4: A fast and versatile genome alignment system. PLoS Comput Biol 14: e1005944. 10.1371/journal.pcbi.1005944.

67. McKenna A, Hanna M, Banks E, Sivachenko A, Cibulskis K, Kernytsky A, Garimella K, Altshuler D, Gabriel S, Daly M, et al. 2010. The Genome Analysis Toolkit: a MapReduce framework for analyzing next-generation DNA sequencing data. Genome Res 20: 1297–1303. 10.1101/gr.107524.110.

68. Minh BQ, Schmidt HA, Chernomor O, Schrempf D, Woodhams MD, von Haeseler A, Lanfear R. 2020. IQ-TREE 2: New models and efficient methods for phylogenetic inference in the genomic era. Mol Biol Evol 37: 1530–1534. 10.1093/molbev/msaa015.

69. Mohamadi H, Chu J, Coombe L, Warren R, Birol I. 2020. ntHits: de novo repeat identification of genomics data using a streaming approach. *bioRxiv* 2020.11.02.365809. https://www.biorxiv.org/content/10.1101/2020.11.02.365809v1.

70. Murphy WJ, Foley NM, Bredemeyer KR, Gatesy J, Springer MS. 2021. Phylogenomics and the Genetic Architecture of the Placental Mammal Radiation. Annu Rev Anim Biosci 9: 29–53. 10.1146/annurev-animal-061220-023149.

71. Murphy WJ, Frönicke L, O’Brien SJ, Stanyon R. 2003. The origin of human chromosome 1 and its homologs in placental mammals. Genome Res 13: 1880–1888. 10.1101/gr.1022303.

72. O’Brien SJ, Menotti-Raymond M, Murphy WJ, Nash WG, Wienberg J, Stanyon R, Copeland NG, Jenkins NA, Womack JE, Marshall Graves JA. 1999. The promise of comparative genomics in mammals. Science 286: 458–62, 479–81. 10.1126/science.286.5439.458.

73. Osmanski AB, Paulat NS, Korstian J, Grimshaw JR, Halsey M, Sullivan KAM, Moreno-Santillán DD, Crookshanks C, Roberts J, Garcia C, et al. 2023. Insights into mammalian TE diversity through the curation of 248 genome assemblies. Science 380: eabn1430. 10.1126/science.abn1430.

74. Paigen K, Petkov P. 2010. Mammalian recombination hot spots: properties, control and evolution. Nat Rev Genet 11: 221–233. 10.1038/nrg2712.

75. Paigen K, Petkov PM. 2018. PRDM9 and Its Role in Genetic Recombination. Trends Genet 34: 291–300. 10.1016/j.tig.2017.12.017.

76. Palsson G, Hardarson MT, Jonsson H, Steinthorsdottir V, Stefansson OA, Eggertsson HP, Gudjonsson SA, Olason PI, Gylfason A, Masson G, et al. 2025. Complete human recombination maps. Nature 639: 700–707. 10.1038/s41586-024-08450-5.

77. Paradis E, Schliep K. 2019. ape 5.0: an environment for modern phylogenetics and evolutionary analyses in R. Bioinformatics 35: 526–528. 10.1093/bioinformatics/bty633.

78. Parvanov ED, Petkov PM, Paigen K. 2010. Prdm9 controls activation of mammalian recombination hotspots. Science 327: 835. 10.1126/science.1181495.

79. Peñalba JV, Wolf JBW. 2020. From molecules to populations: appreciating and estimating recombination rate variation. Nat Rev Genet 21: 476–492. 10.1038/s41576-020-0240-1.

80. Phillips MJ, Bennett TH, Lee MSY. 2009. Molecules, morphology, and ecology indicate a recent, amphibious ancestry for echidnas. Proc Natl Acad Sci U S A 106: 17089–17094. 10.1073/pnas.0904649106.

81. Pollard KS, Hubisz MJ, Rosenbloom KR, Siepel A. 2010. Detection of nonneutral substitution rates on mammalian phylogenies. Genome Res 20: 110–121. 10.1101/gr.097857.109.

82. Poplin R, Ruano-Rubio V, DePristo MA, Fennell TJ, Carneiro MO, Van der Auwera GA, Kling DE, Gauthier LD, Levy-Moonshine A, Roazen D, et al. 2017. Scaling accurate genetic variant discovery to tens of thousands of samples. *bioRxiv* 201178. https://www.biorxiv.org/content/10.1101/201178v3.abstract.

83. Quigley S, Damas J, Larkin DM, Farré M. 2023. syntenyPlotteR: a user-friendly R package to visualize genome synteny, ideal for both experienced and novice bioinformaticians. Bioinform Adv 3: vbad161. 10.1093/bioadv/vbad161.

84. Quinlan AR, Hall IM. 2010. BEDTools: a flexible suite of utilities for comparing genomic features. Bioinformatics 26: 841–842. https://academic.oup.com/bioinformatics/article-pdf/26/6/841/48854754/bioinformatics_26_6_841.pdf.

85. Raudsepp T, Lee E-J, Kata SR, Brinkmeyer C, Mickelson JR, Skow LC, Womack JE, Chowdhary BP. 2004. Exceptional conservation of horse-human gene order on X chromosome revealed by high-resolution radiation hybrid mapping. Proc Natl Acad Sci U S A 101: 2386–2391. 10.1073/pnas.0308513100.

86. Revell LJ. 2012. phytools: an R package for phylogenetic comparative biology (and other things): phytools: R package. Methods Ecol Evol 3: 217–223. 10.1111/j.2041-210X.2011.00169.x.

87. Rhie A, Walenz BP, Koren S, Phillippy AM. 2020. Merqury: reference-free quality, completeness, and phasing assessment for genome assemblies. Genome Biol 21: 245. 10.1186/s13059-020-02134-9.

88. Richard F, Lombard M, Dutrillaux B. 2003. Reconstruction of the ancestral karyotype of eutherian mammals. Chromosome Res 11: 605–618. 10.1023/a:1024957002755.

89. Ritz KR, Noor MAF, Singh ND. 2017. Variation in Recombination Rate: Adaptive or Not? Trends Genet 33: 364–374. 10.1016/j.tig.2017.03.003.

90. Schumer M, Xu C, Powell DL, Durvasula A, Skov L, Holland C, Blazier JC, Sankararaman S, Andolfatto P, Rosenthal GG, et al. 2018. Natural selection interacts with recombination to shape the evolution of hybrid genomes. Science 360: 656–660. 10.1126/science.aar3684.

91. Segura J, Ferretti L, Ramos-Onsins S, Capilla L, Farré M, Reis F, Oliver-Bonet M, Fernández-Bellón H, Garcia F, Garcia-Caldés M, et al. 2013. Evolution of recombination in eutherian mammals: insights into mechanisms that affect recombination rates and crossover interference. Proc Biol Sci 280: 20131945. 10.1098/rspb.2013.1945.

92. Shumate A, Salzberg SL. 2020. Liftoff: accurate mapping of gene annotations. Bioinformatics 37: 1639–1643. 10.1093/bioinformatics/btaa1016.

93. Smit, AFA, Hubley, R, Green, P. RepeatMasker Open-4.0. 2013-2015 http://www.repeatmasker.org.

94. Steinmetz M, Minard K, Horvath S, McNicholas J, Srelinger J, Wake C, Long E, Mach B, Hood L. 1982. A molecular map of the immune response region from the major histocompatibility complex of the mouse. Nature 300: 35–42. 10.1038/300035a0.

95. Stevison LS, Woerner AE, Kidd JM, Kelley JL, Veeramah KR, McManus KF, Great Ape Genome Project, Bustamante CD, Hammer MF, Wall JD. 2016. The Time Scale of Recombination Rate Evolution in Great Apes. Mol Biol Evol 33: 928–945. 10.1093/molbev/msv331.

96. Svartman M. 2012. Chromosome evolution in Xenarthra: new insights from an ancient group. Cytogenet Genome Res 137: 130–143. 10.1159/000339115.

97. Svartman M, Stone G, Stanyon R. 2006. The ancestral eutherian karyotype is present in Xenarthra. PLoS Genet 2: e109. 10.1371/journal.pgen.0020109.

98. Szklarczyk D, Kirsch R, Koutrouli M, Nastou K, Mehryary F, Hachilif R, Gable AL, Fang T, Doncheva NT, Pyysalo S, et al. 2023. The STRING database in 2023: protein-protein association networks and functional enrichment analyses for any sequenced genome of interest. Nucleic Acids Res 51: D638–D646. 10.1093/nar/gkac1000.

99. Van der Auwera GA, Carneiro MO, Hartl C, Poplin R, Del Angel G, Levy-Moonshine A, Jordan T, Shakir K, Roazen D, Thibault J, et al. 2013. From FastQ data to high confidence variant calls: the Genome Analysis Toolkit best practices pipeline: The genome analysis toolkit best practices pipeline. Curr Protoc Bioinformatics 43: 11.10.1-11.10.33. 10.1002/0471250953.bi1110s43.

100. Vandiedonck C, Knight JC. 2009. The human Major Histocompatibility Complex as a paradigm in genomics research. Brief Funct Genomic Proteomic 8: 379–394. 10.1093/bfgp/elp010.

101. Wang Y, Tang H, Debarry JD, Tan X, Li J, Wang X, Lee T-H, Jin H, Marler B, Guo H, et al. 2012. MCScanX: a toolkit for detection and evolutionary analysis of gene synteny and collinearity. Nucleic Acids Res 40: e49. 10.1093/nar/gkr1293.

102. Warren RL, Coombe L, Mohamadi H, Zhang J, Jaquish B, Isabel N, Jones SJM, Bousquet J, Bohlmann J, Birol I. 2019. ntEdit: scalable genome sequence polishing. Bioinformatics 35: 4430–4432. 10.1093/bioinformatics/btz400.

103. Yang F, Alkalaeva EZ, Perelman PL, Pardini AT, Harrison WR, O’Brien PCM, Fu B, Graphodatsky AS, Ferguson-Smith MA, Robinson TJ. 2003. Reciprocal chromosome painting among human, aardvark, and elephant (superorder Afrotheria) reveals the likely eutherian ancestral karyotype. Proc Natl Acad Sci U S A 100: 1062–1066. 10.1073/pnas.0335540100.

104. Zoonomia Consortium. 2020. A comparative genomics multitool for scientific discovery and conservation. Nature 587: 240–245. https://www.nature.com/articles/s41586-020-2876-6.

