## Supplementary Methods, Tables and Figures for "Comparative Genomics Reveals the Ancestral Recombination Landscape of Placental Mammals"

### SUPPLEMENTAL MATERIAL

This File Contains:

- Supplemental Methods
- Supplemental Tables S6-S8
- Supplemental Figures S1-S9
- Supplementary References

#### Supplementary Methods

*DESCHRAMBLER Sensitivity Analysis*

We changed the outgroup to either the aardvark, the sloth, or both the sloth and the aardvark. In addition to humans, we used the sloth as our reference. When the sloth was the outgroup at 500 Kbp block resolution, APCFs similar to those obtained when the aardvark was the outgroup were produced. When we decreased the block resolution to 300 Kbp from the resolution used in our previous run, more APCFs were produced. Most APCFs were similar to the ancestor generated with a 500 Kbp block resolution. However, human chromosome 17 was split into three APCFs; one APCF contained both human chromosomes 18 and 22, a portion of human chromosome 16 was found in another APCF from the APCF containing 16/7, and X was split into two APCFs. Most APCFs were identified when the sloth and aardvark were used as outgroups. Human chromosomes were divided into multiple APCFs; for example, human chromosome 17 is split into seven APCFs in both block resolutions. The number of APCFs did not decrease when the sloth, instead of the human, was used as the reference. Human chromosomes were found in multiple APCFs, including chromosomes 13 and 17. Overall, the following human chromosome(s) and associations were produced into one APCF regardless of the input DESCHRAMBLER parameter changes: 1, 5, 6, 9, 8q, 19q/16q, 19p, and 20. These chromosome(s)/associations were primarily found in a single APCF except in all but a single run: 3/21, 8p/4, 2q, 7a, 2pq, 11, 13, 10q, 18, and X. The human chromosome(s) or associations 14/15, 10p/12pq/22qt, 17, 7b/16p, and 12qt/22q were found in multiple APCFs.

**Table S6.** DESCHRAMBLER Sensitivity Analysis

| **Reference and Outgroup Used** | **Human reference; Aardvark Outgroup** | **Human reference; Sloth Outgroup** | **Human reference; Sloth and Aardvark Outgroup** | **Sloth reference; Aardvark Outgroup** |
|---|---|---|---|---|
| **Number of ancestral fragments that agree with Svartman (2012)** | 18 | 18 | 11 | 16 |

**Supplemental Table S7.** Genome Assemblies Used in this Study

| **Species** | **Common name** | **NCBI accession number** | **Source** |
| --- | --- | --- | --- |
| *Homo sapiens* | Human | GCF_000001405.40<br>GCF_009914755.1 (Aardvark and sloth alignments) | PRJNA31257<br>Nurk et al. 2022 (Aardvark and sloth alignments) |
| *Felis catus* | Domestic Cat | GCF_018350175.1 | PRJNA684600 |
| *Orycteropus afer* | Aardvark | PRJNA1073880<br>GCF_000298275.1 (Liftoff) | **This study**<br>PRJNA74587 (Liftoff) |
| *Choloepus hoffmanni* | Hoffmann's two-toed sloth | PRJNA1073880 | **This study** |
| *Balaenoptera musculus* | Blue Whale | GCF_009873245.2 | Bukhman et al. 2024 |
| *Choloepus didactylus* | Linnaeus's two-toed sloth | GCF_015220235.1 | PRJNA561937 |
| *Bos taurus* | Domestic cattle | GCF_002263795.3 | PRJNA391427 |
| *Dasypus novemcinctus* | Nine-banded armadillo | GCF_030445035.2<br>GCF_030445035.1 (Liftoff) | PRJNA923800 |
| *Myotis myotis* | Greater mouse-eared bat | GCF_014108235.1 | PRJNA628559 |
| *Sus scrofa* | Pig | GCF_000003025.6 | Warr et al. 2020 |
| *Elephas maximus* | Asian Elephant | GCF_024166365.1 | PRJNA850184 |
| *Canis lupus familiaris* | Domestic Dog | GCF_000002285.5 | Jagannathan et al. 2021 |
| *Mus musculus* | House Mouse | GCF_000001635.27 | PRJNA20689 |
| *Ceratotherium simum* | White Rhino | GCA_023653735.1<br>GCF_000283155.1 (Liftoff) | PRJNA786211<br>PRJNA74583 (Liftoff) |

**Supplemental Table S8.** Software and Algorithms Used in this Study

| Program | Version | Parameters |
|---|---|---|
| Flye | 2.8.3-b1695 | --pacbio-raw |
| NextDenovo | 2.2-beta.0 | Default except seed_cutoff=15470; blocksize=1g; pa_raw_align=4; pa_correction=4; sort_options = -m 1g -t 4 -k 50; minimap2_options_raw = -x ava-pb -t 10; correction_options = -p15; random_round = 100; minimap2_options_cns = -x ava-ont -t 10 -k17 -w17 |
| Juicer | 1.6 | -s MboI |
| 3D-DNA | 180419 | OAF[1]: --input 1000 –rounds 3<br>CHO[2]: --input 15000 –rounds 1 |
| Juicebox Assembly Tools | 1.11.08 | N/A |
| FCA-adaptor | 0.2.3 | --euk |
| Mummer | 4.0.0rc1 | Nucmer default settings |
| ntHits | 0.1.0 | --solid –outbloom -b36 -k 50 |
| ntedit | 1.3.5 | Default parameters |
| Hapo-G | 1.3.4 | Default parameters: 2 rounds |
| Burrows-Wheeler Aligner (BWA) | 0.7.17 | Default mem parameters |
| Samtools | 1.13 | samtools view -F 0x4 -b - \| samtools fixmate -m - \| samtools sort -m 2g - \| samtools markdup -r - |
| Meryl | 1.3 | k=21 |
| Merqury | 1.3 | Default parameters |
| Compleasm | 0.2.6 | Default parameters; mammalia_odb10 library used |
| QUAST | 5.3.0 | Default parameters |
| Minimap2 | 2.23 | -ax map-pb -I10g |
| Samtools | 1.12 | samtools view -bh - \| samtools sort - |
| Geneious Prime | 2022.1.1 | N/A |
| BLAST+ | 2.12.0 | -task dc-megablast -use_index true |

[1]OAF = aardvark [2]CHO = Hoffmann's two-toed sloth

**Supplemental Table S8.** Software and Algorithms Used in this Study Continued

| Program | Version | Parameters |
|---|---|---|
| GENESPACE (gene annotations) | 1.3.1 | Default Parameters |
| MCScanX | 2022.10.31 | Default Parameters |
| DIAMOND | 2.0.15 | Default Parameters |
| OrthoFinder | 2.5.4 | Default Parameters |
| Liftoff | 1.6.3 | OAF[1], DNO[3] and CSI[4]: Default parameters<br>CHO[2]: -flank 0.5 -d 3 |
| Another Gtf/Gff Analysis Toolkit (AGAT) | 0.9.2 | agat_sp_extract_sequences.pl -p |
| GENESPACE (whole genome alignments) | 1.4.1 | daisyChain= TRUE; minMapq = 60, keepNSecondary = 0; windowSize = 5000; minChrSize = 500e3 |
| MCScanX | 2022.10.31 | Default Parameters |
| DIAMOND | 2.1.0 | Default Parameters |
| Minimap2 | 2.24 | Default Parameters |
| Lastz_32 | 1.04.15 | --notransition --step=20 --nogapped --format=axt –ambiguous=iupac |
| GenomeAlignmentTools | 2019.11.20 | axtChain -linearGap=medium -minScore=1000; chainSplit (default); chainSort (default); chainPreNet (default); chainNet (default) |
| DESCHRAMBLER | N/A | Our ancestor:<br>REFSPC = Human RESOLUTION = 500000 MINADJSCR = 0.0001<br>Outgroup = aardvark<br>Sensitivity analysis:<br>REFSPC = Human or Sloth<br>RESOLUTION = 300000 or 500000<br>Outgroup = aardvark, sloth, or aardvark and sloth |
| BEDtools | 2.30.0 | makewindows -w 1500000<br>human: map -c4 -o mean<br>all other mammals: map -c5 -o mean |

[1]OAF = aardvark [2]CHO = Hoffmann's two-toed sloth [3]DNO = nine-banded armadillo [4]CSI = white rhino

**Supplemental Table S8.** Software and Algorithms Used in this Study Continued

| Program | Version | Parameters |
|---|---|---|
| Mummer | 4.0.0rc1 | show-coords -r -T |
| WebGestalt | N/A | Multiple Test Adjustment = BH, Significance Level = FDR 0.05, Redundancy Removal = Weighted set cover |
| STRING | 12.0 | Default,<br>Redundancy removal = similarity >= 0.5 |

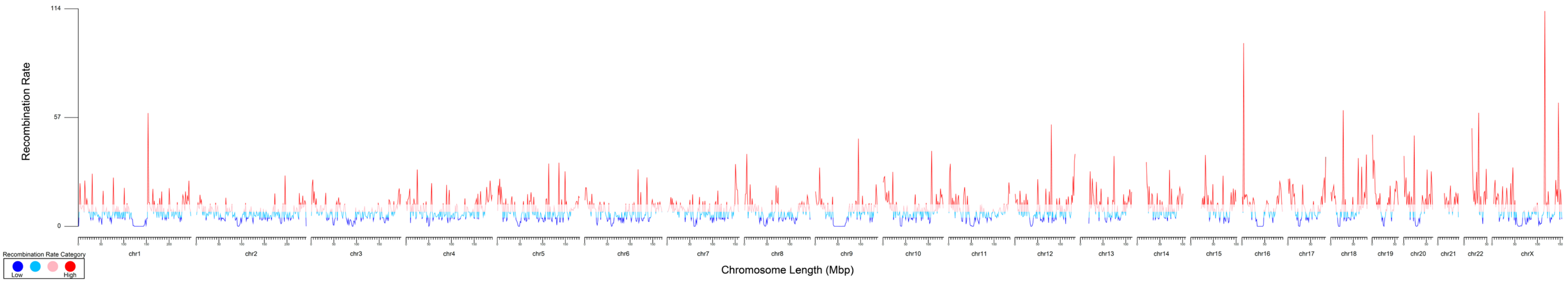


**Supplementary Figure 1.** Recombination map of the human genome. Averaged recombination rates (y-axis) in 1.5 Mb windows are depicted across each chromosome (x-axis). Colors highlight one of the four recombination rate categories, with dark blue being the lowest recombination rate and red being the highest.

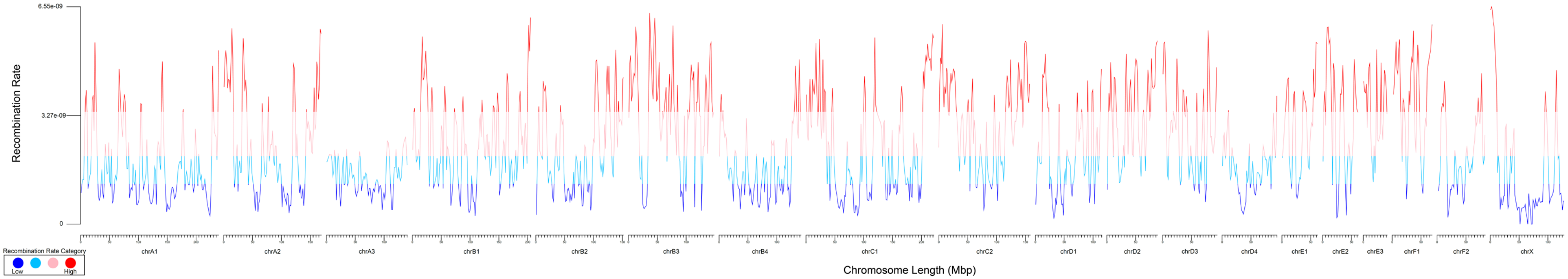


**Supplementary Figure 2.** Recombination map of the domestic cat genome. Averaged recombination rates (y-axis) in 1.5 Mb windows are depicted across each chromosome (x-axis). Colors highlight one of the four recombination rate categories, with dark blue being the lowest recombination rate and red being the highest.

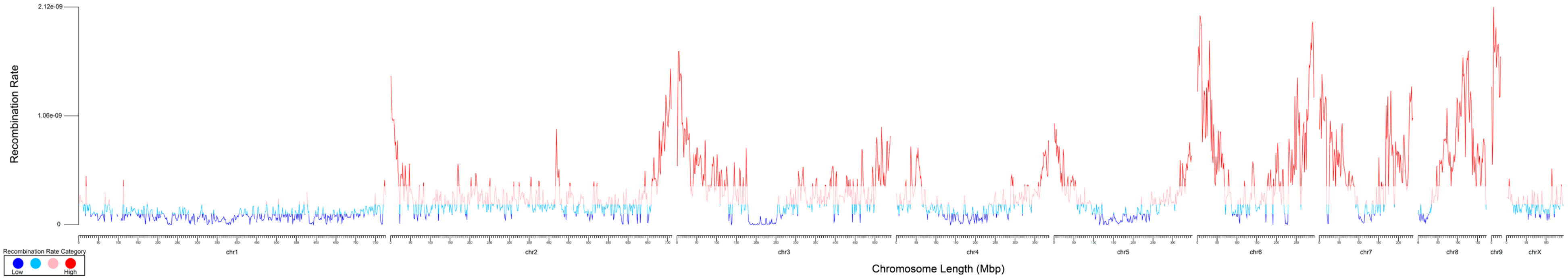


**Supplementary Figure 3.** Recombination map of the aardvark genome. Averaged recombination rates (y-axis) in 1.5 Mb windows are depicted across each chromosome (x-axis). Colors highlight one of the four recombination rate categories, with dark blue being the lowest recombination rate and red being the highest.

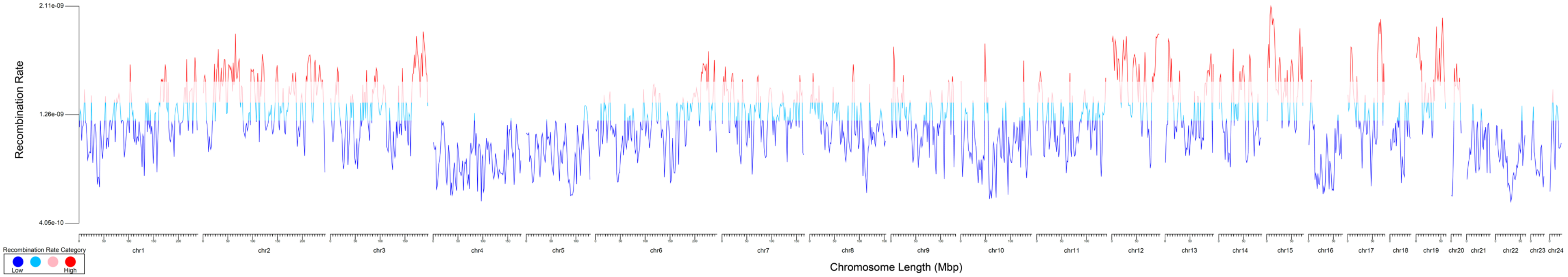


**Supplementary Figure 4.** Recombination map of the Hoffmann's two-toed sloth genome. Averaged recombination rates (y-axis) in 1.5 Mb windows are depicted across each chromosome (x-axis). Colors highlight the four recombination rate categories, with dark blue representing the lowest and red the highest.

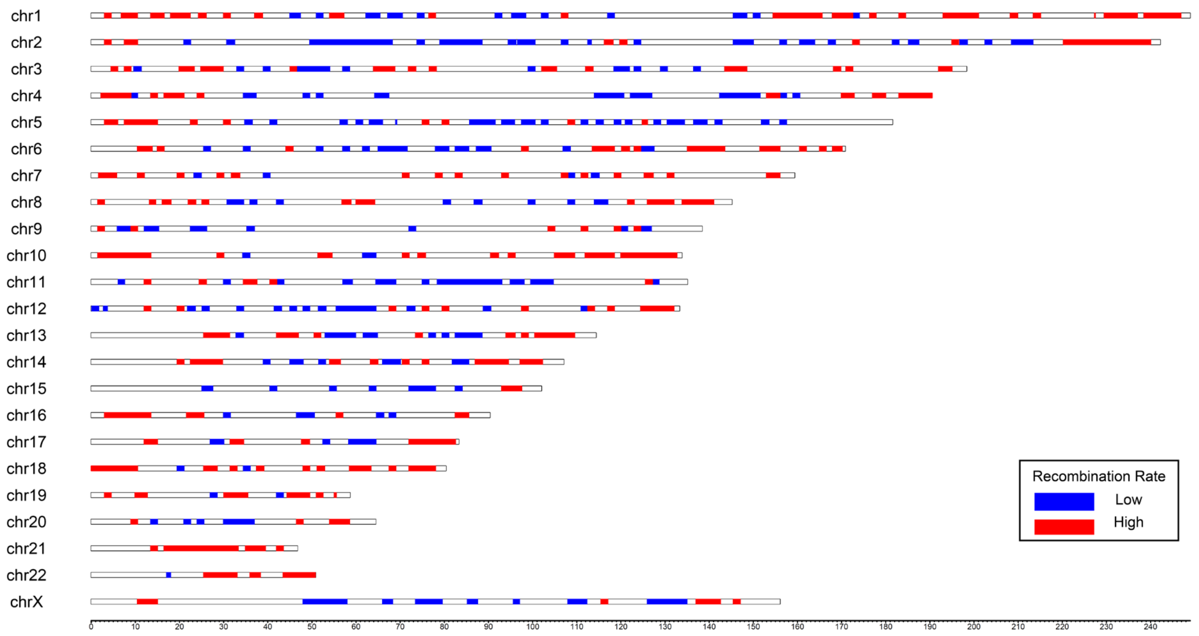


**Supplementary Figure 5.** Human ideogram illustrates syntenic regions with low and high meiotic recombination rates. Colors along the chromosomes represent the conserved regions, with blue indicating low and red indicating high recombination rates. The x-axis depicts the length of the chromosomes in base pairs.

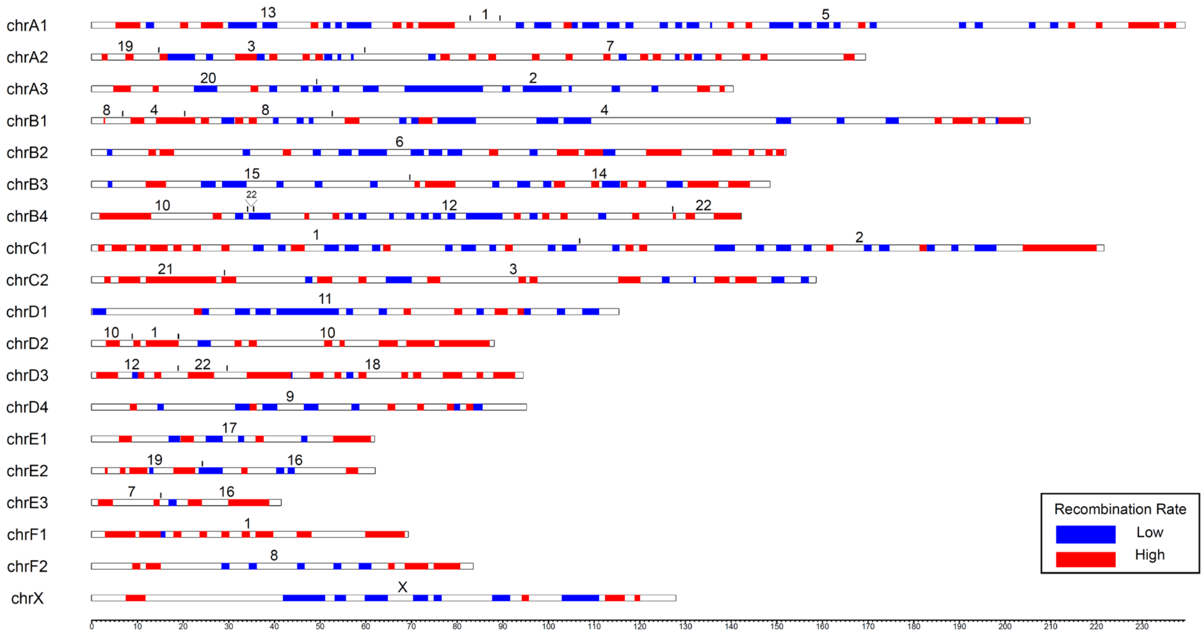


**Supplementary Figure 6.** Domestic cat ideogram illustrates syntenic regions with low and high meiotic recombination rates. Colors along the chromosomes represent the conserved regions, with blue indicating low and red indicating high recombination rates. The numbers above each chromosome indicate the corresponding human chromosome. The x-axis depicts the length of the chromosomes in base pairs.

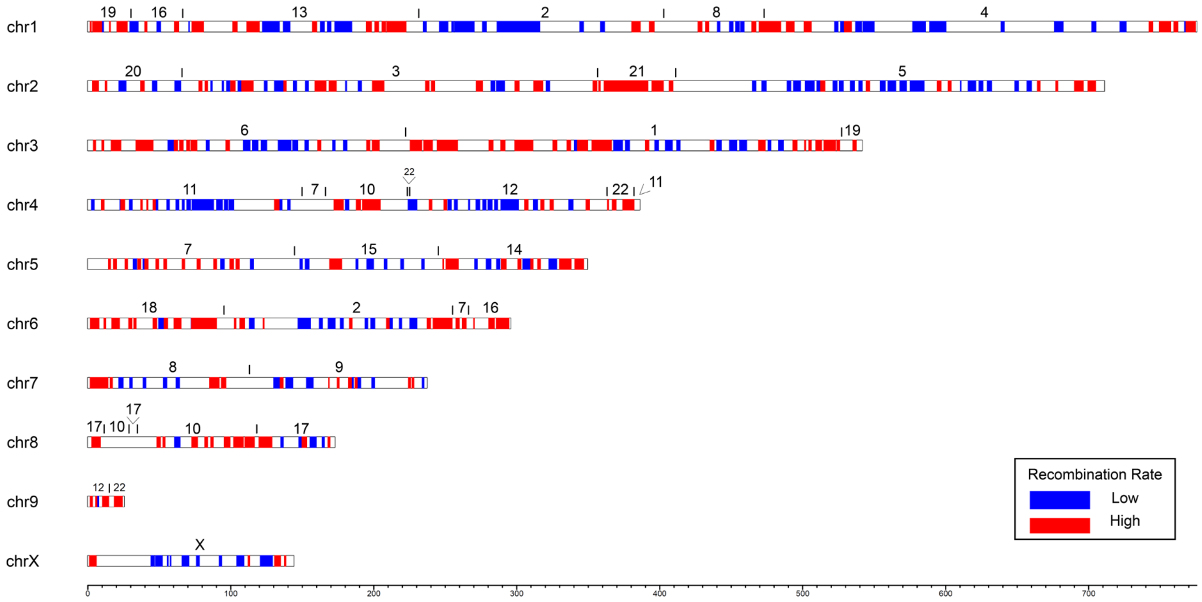


**Supplementary Figure 7.** Aardvark ideogram illustrates syntenic regions with low and high meiotic recombination rates. Colors along the chromosomes represent the conserved regions, with blue indicating low and red indicating high recombination rates. The numbers above each chromosome indicate the corresponding human chromosome. The x-axis depicts the length of the chromosomes in base pairs.

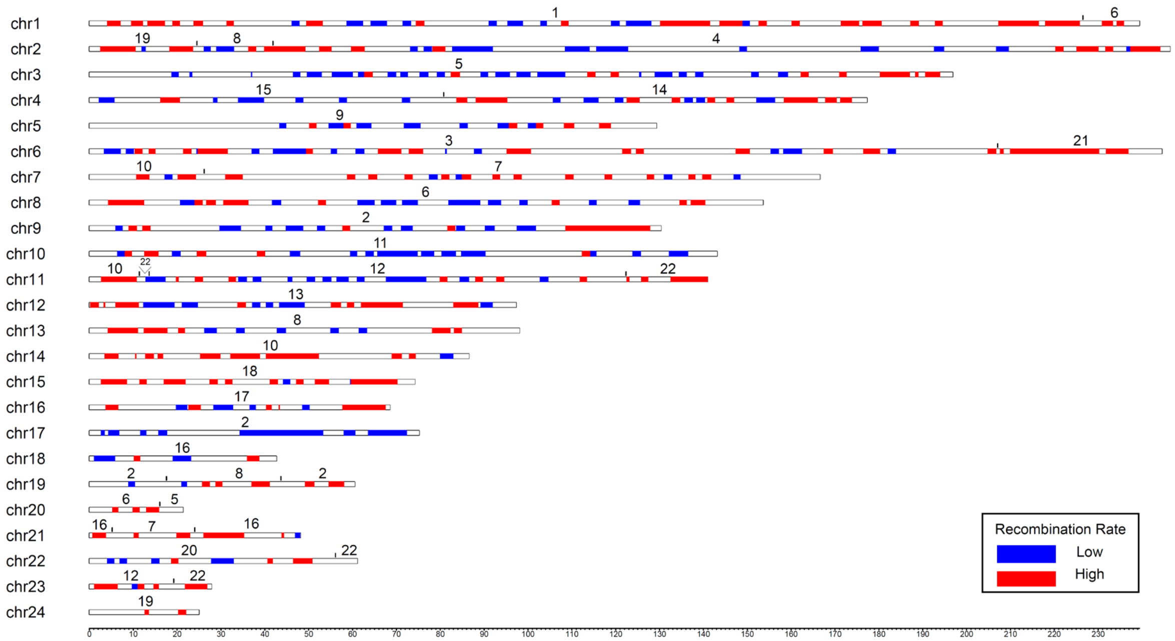


**Supplementary Figure 8.** Hoffmann's two-toed sloth ideogram illustrates syntenic regions with low and high meiotic recombination rates. Colors along the chromosomes represent the conserved regions, with blue indicating low and red indicating high recombination rates. The numbers above each chromosome indicate the corresponding human chromosome. The x-axis depicts the length of the chromosomes in base pairs.

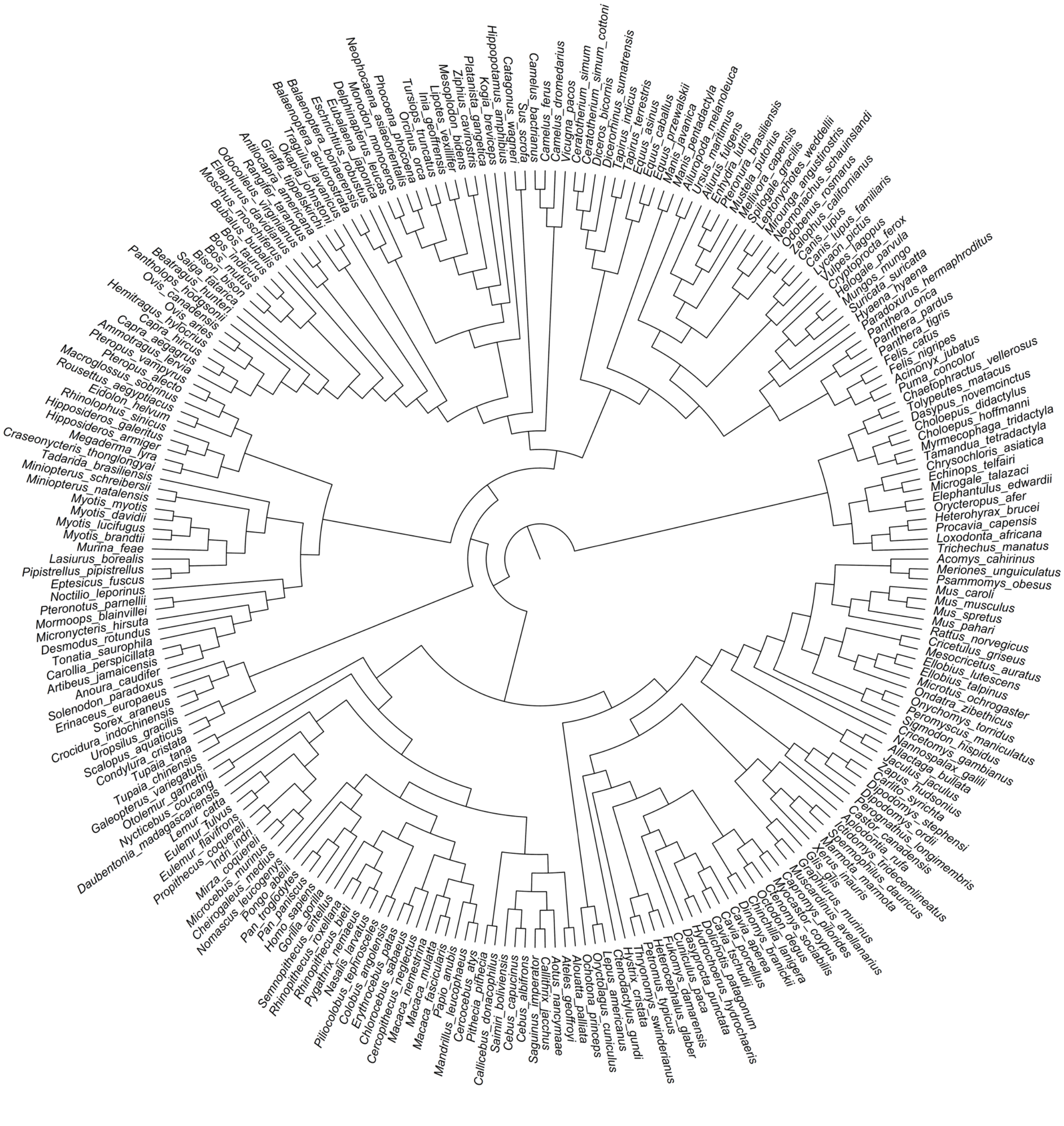

**Supplementary Figure 9.** Rooted tree from the regions of low recombination from the human-referenced Zoonomia alignment (Foley et al. 2023).

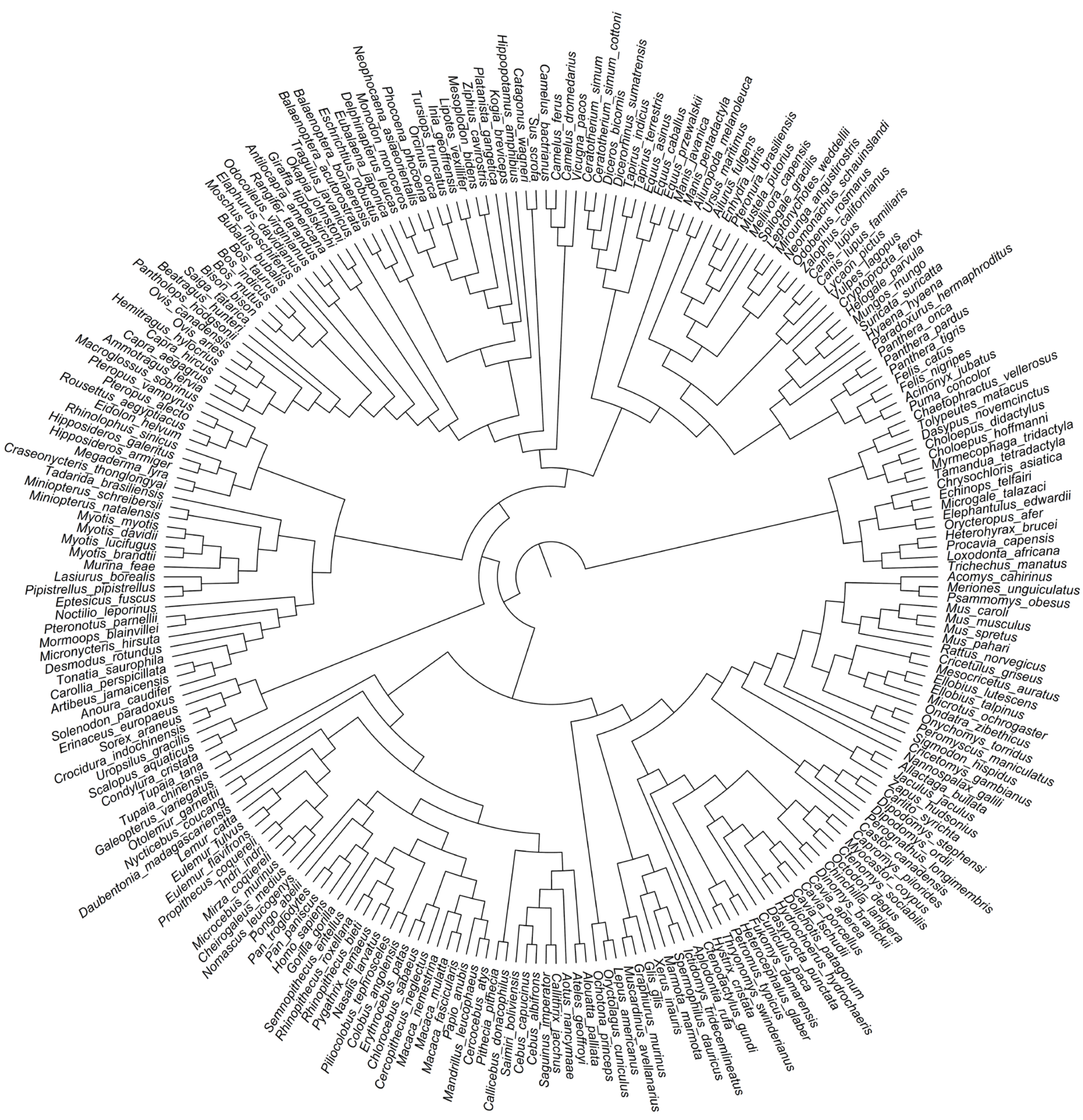


**Supplementary Figure 10.** Rooted tree from the regions of high recombination from the human-referenced Zoonomia alignment (Foley et al. 2023).

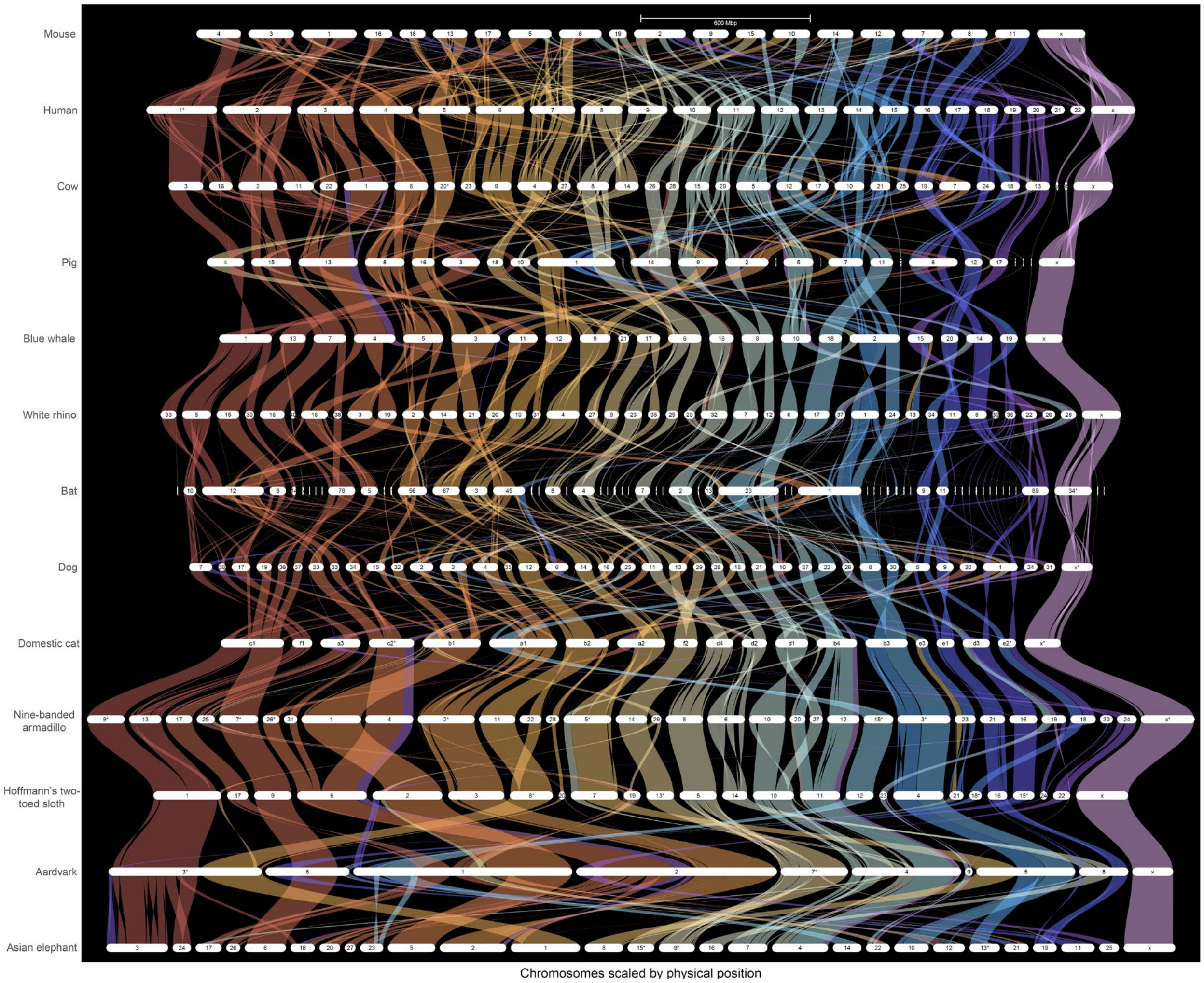


**Supplementary Figure 11.** Gene synteny plot generated by GENESPACE for thirteen divergent mammals. Solid bands are regions of collinearity shared between the thirteen species. Colored bands connecting chromosomes indicate conserved synteny relative to the human.

Supplementary References

Bukhman YV, Morin PA, Meyer S, Chu L-F, Jacobsen JK, Antosiewicz-Bourget J, Mamott D, Gonzales M, Argus C, Bolin J, et al. 2024. A high-quality blue whale genome, segmental duplications, and historical demography. *Mol Biol Evol* **41**. http://dx.doi.org/10.1093/molbev/msae036.

Jagannathan V, Hitte C, Kidd JM, Masterson P, Murphy TD, Emery S, Davis B, Buckley RM, Liu Y-H, Zhang X-Q, et al. 2021. Dog10K_boxer_Tasha_1.0: A long-read assembly of the dog reference genome. *Genes (Basel)* **12**: 847. http://dx.doi.org/10.3390/genes12060847.

Nurk S, Koren S, Rhie A, Rautiainen M, Bzikadze AV, Mikheenko A, Vollger MR, Altemose N, Uralsky L, Gershman A, et al. 2022. The complete sequence of a human genome. *Science* **376**: 44–53. http://dx.doi.org/10.1126/science.abj6987.

Warr A, Affara N, Aken B, Beiki H, Bickhart DM, Billis K, Chow W, Eory L, Finlayson HA, Flicek P, et al. 2020. An improved pig reference genome sequence to enable pig genetics and genomics research. *Gigascience* **9**: giaa051. http://dx.doi.org/10.1093/gigascience/giaa051.
